# Structural Basis of Human Kinetochore-Microtubule Coupling by the Ndc80 and Ska Complexes

**DOI:** 10.64898/2025.12.06.692780

**Authors:** Ju Zhou, Ipsita Saha, Valerie Siahaan, Yuanchang Zhao, Carsten Janke, Ahmet Yildiz, Eva Nogales

**Author notes:** Correspondence (A.Y.), (E.N.). These authors contributed equally to this work (Ju Zhou, Ipsita Saha).

## Abstract

Faithful chromosome segregation depends on kinetochore-microtubule attachments that must be load-bearing, dynamic, and tightly regulated. In human cells, the Ska and Ndc80 complexes (SkaC and Ndc80C) cooperate to sustain these attachments, but how they are organized on microtubules and coupled to end dynamics remains unclear. Here, our cryo-EM structure shows that Ndc80C and SkaC form a coupled assembly across neighboring protofilaments. A phosphorylation-sensitive SKA3 tether binds a defined kink in the Ndc80 coiled-coil, positioning SKA1 beside Ndc80C. These two complexes jointly sandwich the α-tubulin tail and sense its polyglutamylation state. SkaC recognizes the extended microtubule lattice and, through the SKA3 tether, confers this selectivity onto Ndc80C, enabling Ndc80C tip tracking while strengthening microtubule stabilization. Together, our findings define the structural basis of Ndc80C-SkaC cooperation and uncover a phosphorylation-regulated, polyglutamylation-sensitive, and lattice-state-dependent mechanism for robust and regulated KT-MT coupling during chromosome segregation in mitosis.

## INTRODUCTION

During cell division, accurate segregation of chromosomes requires kinetochore-microtubule (KT-MT) attachments that are both mechanically robust and dynamically adaptable^1-3^. These attachments must sustain force while remaining coupled to MT plus ends as they grow and shorten during chromosome congression and segregation^4-6^. Failure to maintain this balance can produce erroneous or hyperstable attachments that evade error correction, leading to chromosome mis-segregation, aneuploidy, and chromosome instability associated with cancer and developmental disease^7-10^.

At the outer KT, the KNL1, Mis12, and Ndc80 complexes form the conserved KMN network that links chromosomes to spindle MTs^2,11^. The Ndc80C is the principal MT-binding and force-coupling element of this network^4,5^. Human Ndc80C is an elongated heterotetramer composed of NDC80, NUF2, SPC24, and SPC25, with the NDC80-NUF2 calponin-homology domains binding MTs and the SPC24-SPC25 end connecting to inner KT receptors^12-16^. Although Ndc80C can bind cooperatively along MTs, how this core attachment complex is converted into a regulated assembly that remains coupled to dynamic MT plus ends remains unresolved^2,5,14^.

Eukaryotic KTs share the challenge of maintaining persistent attachment to dynamic MT ends, but the molecular solutions to this problem have diverged. In budding yeast, Dam1C forms oligomeric MT-associated assemblies and couples to Ndc80C through defined interfaces^17-22^. Metazoan KTs lack Dam1C and instead rely on SkaC to reinforce mature KT-MT attachments^23-25^. SkaC accumulates at KTs as MT attachments mature^23,26^, promotes stable end-on attachment^27,28^, prevents force-dependent detachment^26^, and supports MT-end tracking^24,29-31^. SkaC is organized as a W-shaped homodimer of SKA1-SKA2-SKA3 heterotrimers^32,33^. SKA1 contains an MT-binding domain (MTBD), whereas the extended C-terminal region of SKA3 interacts with Ndc80C in a manner enhanced by cyclin-dependent kinase phosphorylation^30,31,33,34^. While these properties establish SkaC as a key regulator of mature KT-MT, the molecular architecture by which SkaC is physically coupled to Ndc80C on the MT surface, and how this coupling contributes to MT dynamics and supports persistent plus-end tracking, has remained incompletely defined^30,31,35^.

How SkaC is positioned with respect to Ndc80C on MTs remains unresolved^24-26,30,31,35^. Previous studies have implicated different Ndc80C regions in SkaC binding, including the NDC80 N-terminal tail^35,36^, the internal loop^28,34,37^, and coiled-coil regions^30,33^. Previous studies also proposed distinct modes of SkaC-MT engagement, including single-PF binding^35^, multi-PF or multi-orientation binding^32,33^, and larger oligomeric or multivalent assembly^38^. Therefore, the molecular architecture that organizes human Ndc80C and SkaC into a regulated, plus-end-coupled attachment is still unclear.

By integrating cryo-electron microscopy (cryo-EM) with biochemical assays and single-molecule measurements on dynamic MTs, we define the architecture of the human Ndc80C-SkaC assembly on the MT surface and assess how this organization controls PTM-sensitive MT engagement, lattice-state selectivity, regulation of MT dynamics, and plus-end coupling. Our findings reveal that SkaC is not simply an accessory MT binder, but a regulated partner that, through a SKA3-mediated interface, couples Ndc80C to the structural state and dynamics of MT ends. This mechanism provides a framework for how human KTs maintain robust yet adaptable chromosome attachments.

## RESULTS

### Ndc80C and SkaC simultaneously bind adjacent protofilaments

We purified recombinant full-length human SkaC and Ndc80C for cryo-EM analysis (Extended Data Fig. 1) and used a phosphomimetic SkaC-6D carrying aspartate substitutions in SKA3 (T203D, T217D, S283D, T291D, T358D, and T360D) to mimic phosphorylation that has been shown to enhance interaction with Ndc80C^39^ (Extended Data Fig. 1a). In samples containing Ndc80C alone, MTs were seen uniformly coated by Ndc80C, with sharp edges corresponding to the coiled-coil region parallel to the MT axis of ∼660 Å outer diameter (Extended Data Fig. 2a). In the presence of SkaC-6D, the Ndc80C coiled-coil regions appeared less ordered, indicating a distinct MT decoration pattern (Extended Data Fig. 2b).

We determined the average structure of Ndc80C and SkaC co-bound to MTs at resolutions ranging from 2.8 Å (tubulin) to 6 Å (higher-radius, visible segment of Ndc80C) (Extended Data Figs. 2-4; Supplementary Video 1; Supplementary Table 1). Ordered segments of the complexes span a diameter of approximately 550 Å (Extended Data Fig. 5a), whereas density beyond was not resolved, reflecting its conformational flexibility.

Consistent with prior observations^14,40^, Ndc80C binds along a PF with tubulin-monomer periodicity (Extended Data Fig. 2c,d). By contrast, SkaC binds with a tubulin dimer repeat at the longitudinal interdimer interface between adjacent PFs (Extended Data Figs. 2d and 5a,b). In the averaged structure, the SKA1-MTBD density partially overlaps with that of the NDC80-CHD (Extended Data Fig. 5a,b and Supplementary Video 1), revealing a steric clash and indicating that particles with different Ndc80C- or SkaC-bound registries were mixed during averaging. 3D classification at the single tubulin dimer level confirmed that Ndc80C and SkaC do not simultaneously bind the same tubulin (Extended Data Fig. 2d, left panel).

We next asked whether SkaC and Ndc80C can bind adjacent tubulin dimers along the same PF. 3D classification on two longitudinal dimers identified either no complex bound, Ndc80C bound consecutively to four tubulin monomers, or SkaC bound at the interdimer junction (Extended Data Fig. 2d, middle panel). Thus, SkaC and Ndc80C are unlikely to bind to longitudinally adjacent tubulin dimers, probably due to cooperative Ndc80C binding along a PF. In contrast, classification of two lateral dimers revealed a class in which SkaC and Ndc80C are simultaneously engaging two adjacent PFs (Extended Data Fig. 2d, right panel, and Extended Data Fig. 2e). This result indicates that at the human outer KT, Ndc80C clustering along one PF promotes SkaC recruitment to the adjacent PF (Fig. 1a,b).

**Fig. 1:**
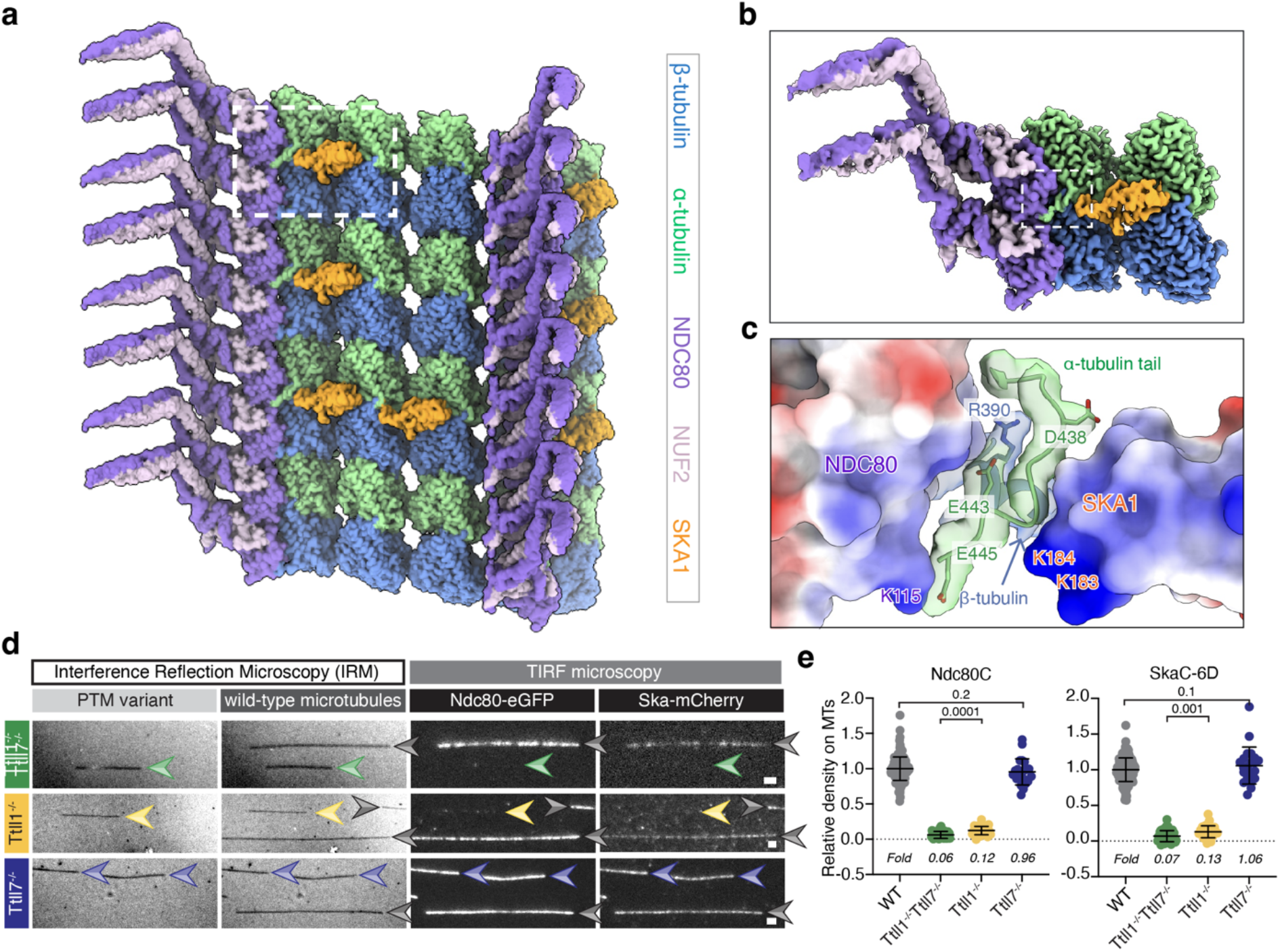
Structural and biochemical basis of the MT-bound Ndc80C-SkaC “sandwich” conformation. a, Composite cryo-EM map illustrating the different arrangements of Ndc80C and SkaC on the MT surface identified by 3D classification. The white dashed box highlights that Ndc80C and SkaC are simultaneously bound to two adjacent PFs. Maps were sharpened using DeepEMhancer for visualization. **b**, Enlarged view of the boxed region in (**a**). **c**, The “sandwich” interface formed by the NDC80 CHD, the α-CTT, and the SKA1-MTBD. NDC80 and SKA1 are shown as electrostatic surfaces. Key residues involved in α-CTT stabilization are indicated. A segment of β-tubulin is also shown, with the R390 side chain displayed as sticks. **d**, GMPCPP MTs assembled from knockout (colored arrowheads) or wild-type (WT) mouse brain tubulin (grey arrowheads) were sequentially immobilized in the chamber and visualized using IRM. 4 nM Ndc80C-eGFP and 4 nM SkaC-mCherry were added to the chamber and imaged using TIRF microscopy. Scale bars: 2 µm. **e**, Quantitative analysis of relative Ndc80C (left) or SkaC (right) density on WT and PTM-variant MTs (mean ± SD). Fluorescence intensities of Ndc80C or SkaC on PTM-variant MTs were normalized to those on WT MTs(n = 169, 44, 37, 22 MTs in 3 independent experiments, respectively). P-values were determined using a two-tailed t-test.

### Ndc80C and SkaC sandwich the α-tubulin tail

Our analysis revealed a previously uncharacterized mode of cooperative tubulin engagement of the two KT complexes: the C-terminal tail of α-tubulin (α-CTT, residues 438-445) is “sandwiched” between the NDC80 CHD and the SKA1 MTBD across two adjacent PFs (Fig. 1b,c). Tubulin tails are negatively charged peptides that often cannot be resolved by cryo-EM due to their inherent flexibility and sequence variation among α-tubulin isoforms. However, α-CTT becomes ordered when sandwiched between SkaC and Ndc80C, and D438, E443, and E445 are conserved (Extended Data Fig. 5c), enabling confident modeling of the α-CTT density up to E445 (Fig. 1c).

The NDC80 and SKA1 interfaces that engage the αCTT are strongly positively charged (Fig. 1c). NDC80 K115 and β-tubulin R390 form salt bridges with α-tubulin E445 and E443, respectively. On the other side of the interface, the α-CTT is engaged by SKA1 K183 and K184. Our structure indicates that αCTT stabilizes the Ndc80C-SkaC interaction by mediating simultaneous interactions with both complexes. This is consistent with the NDC80 K115E mutation to impair KT-MT attachments and chromosome alignment^41^. The interactions we observed between SKA1 and α-CTT also provide an explanation for how SKA1 mutants that neutralize K183 and K184 weaken MT end tracking and increase chromosome alignment defects *in vivo*^42^ while reducing MT binding *in vitro*^33^.

When adjacent PFs are not co-occupied with Ndc80C on the left and SkaC on the right (when facing towards the MT plus-end), the α-CTT extends outward to interact with the N-terminal tail (NTT) of NDC80 in the adjacent PF (Extended Data Fig. 5d), as previously described^14,40^. We also observed stabilization of the β-tubulin CTT, up to residue E435, by interaction with the NDC80 CHD (Extended Data Fig. 5e). These findings further suggest that tubulin isotype-specific CTT sequences and their PTMs can regulate KT-MT interactions.

### Tubulin tails and polyglutamylation modulate cooperative Ndc80C-SkaC binding

Previous studies have shown that Ndc80C binding to MTs depends strongly on the tubulin CTTs^12,33^, whereas the reported effects on SkaC binding have been less consistent^33,43^. To quantify the functional significance of the tubulin tails in Ndc80C and SkaC binding to MTs, we removed the tubulin CTTs by subtilisin treatment and assessed MT binding by pelleting assays. Subtilisin treatment markedly reduced binding of Ndc80C alone, SkaC alone, or co-binding of the two complexes together (Extended Data Fig. 5f). These results indicate that the tubulin tails contribute to both the individual association of Ndc80C and SkaC with MTs^12,43^ and their co-engagement with the MT surface.

α-tubulin E445, the most C-terminal residue seen in our structure, is a preferential site of polyglutamylation, a posttranslational modification (PTM) that regulates MT dynamics and chromosome segregation^44-47^ (Fig. 1c). To test whether tubulin PTMs could directly modulate Ndc80C and SkaC binding, we first compared their binding to HeLa-MTs, which carry low levels of tubulin tail modifications^48^, and porcine brain MTs, which are highly post-translationally modified^49-51^. In MT pelleting assays, Ndc80C alone, SkaC alone, and the two complexes together, all exhibited weaker binding to HeLa MTs than to porcine brain MTs (Extended Data Fig. 6a,b). Consistently, TIRF-based titration experiments showed reduced recruitment of Ndc80C and SkaC to HeLa MTs, particularly at lower concentrations of these complexes (Extended Data Fig. 6c,d). These results are consistent with a role for tubulin PTMs, although differences in tubulin isotype composition may also affect binding (Extended Data Fig. 7a,b).

To assess the contribution of tubulin PTMs more directly, we next focused on polyglutamylation, as the modification that should impart the largest effect on MT binding by both Ndc80C and SkaC based on our structure. Using MTs from mutant mouse brain lacking defined tubulin-polyglutamylation enzymes, we found that absence of both TTLL1 and TTLL7, or TTLL1 alone, strongly reduced binding to MTs by both Ndc80C and SkaC. In contrast, MTs from TTLL7-KO mice retained near wild-type levels of Ndc80C and SkaC binding (Fig. 1f,g). Because TTLL1 preferentially modifies α-tubulin, whereas TTLL7 primarily modifies β-tubulin^52-55^, these results indicate that α-tubulin polyglutamylation promotes Ndc80C-SkaC engagement with MTs.

Thus, our data identify the α-CTT as a key element in an interaction hub for the human outer KT that can be regulated via polyglutamylation. Sandwiching of the α-CTT between Ndc80C and SKA1 provides a molecular explanation for how tubulin-tail modifications, particularly α-tubulin polyglutamylation, can tune the cooperative recruitment of Ndc80C and SkaC to the MT lattice.

### Molecular interactions at the MT-Ndc80C interface

As previously shown at lower resolution^14,40^, alternating Ndc80Cs contact α- and β-tubulin along a PF (Fig. 2a). We denote the complexes that predominantly engage α-tubulin or β-tubulin as α-Ndc80C and β-Ndc80C, respectively (Fig. 2a). The α-Ndc80C interface with α-tubulin involves a combination of electrostatic, hydrogen-bonding, and hydrophobic contacts (Fig. 2b,c and Extended Data Fig. 8a-c). In particular, residues K115, K166, S165/S167, and H176 contribute to α-tubulin engagement, with NDC80 S165 notable as a reported NEK2 phosphorylation site^56-58^ (Fig. 2b and Supplementary Video 1). Additional hydrophobic contacts centered on A174, P175, and H176 further reinforce this interface (Fig. 2b and Extended Data Fig. 8c). By contrast, α-Ndc80C makes fewer contacts with β-tubulin (Fig. 2c and Extended Data Fig. 8d). β-Ndc80C engages tubulin in a closely analogous manner, consistent with conservation of the relevant contact surfaces in α- and β-tubulin (Extended Data Fig. 8e,f) and with the monomer-repeat binding pattern of Ndc80C on MTs.

**Fig. 2:**
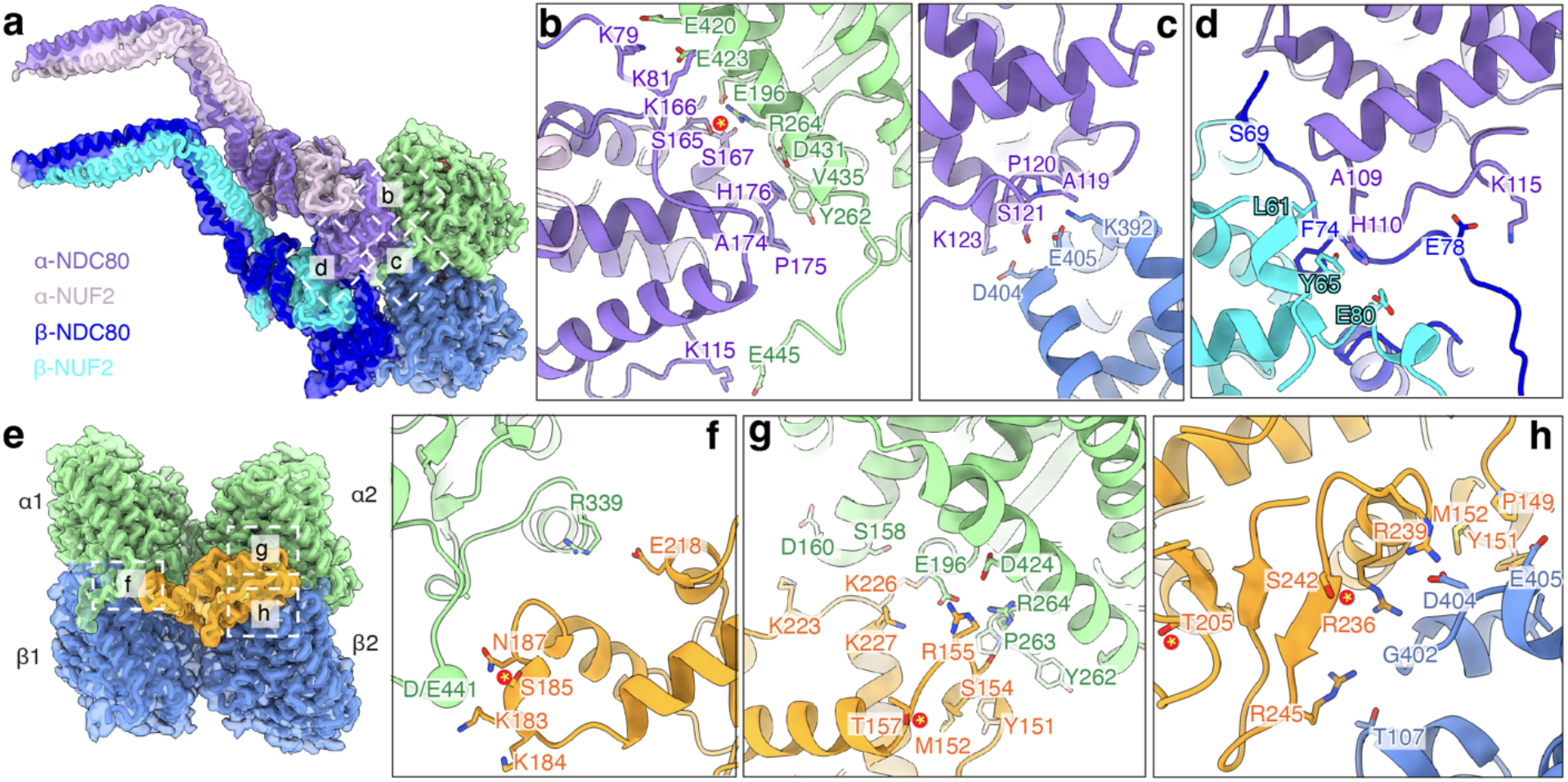
Distinct MT interactions shape binding of Ndc80C and SkaC to MTs. a, Cryo-EM map with fitted models showing two adjacent Ndc80Cs, denoted as α-Ndc80C and β-Ndc80C, bound to a tubulin dimer and to each other. Regions shown in detail on the right are indicated. Maps were sharpened using DeepEMhancer for visualization (same for (**e**)). **b**, Close-up view of the interface between α-Ndc80C and α-tubulin. Key residues are labeled and shown as sticks, and phosphorylation sites are marked by asterisks (same for panels (**c**-**h**)). **c**, Close-up view of the interface between α-Ndc80C and β-tubulin. **d**, Interface between two adjacent Ndc80C. **e**, Cryo-EM map with fitted models showing the SKA1-MTBD (orange) bound at the interdimer interface between two adjacent PFs. For clarity, tubulin subunits are labeled α1, α2, β1, and β2. The regions shown in detail on the right are indicated. **f**, Close-up view of the interface between SKA1 and α1-tubulin. **g**, Close-up view of the interface between SKA1 and α2-tubulin. **h**, Close-up view of the interface between SKA1 and β2-tubulin.

### Cooperative binding of Ndc80C on the MT surface

As human Ndc80C binds to both tubulin monomers (Fig. 2a), it self-associates into clusters along a PF^14,40^. Direct inter-complex contacts are mediated in part by the NDC80 NTT, with β-NDC80 residues N68-K81 inserting into a groove formed between α-NDC80 and β-NUF2 (Fig. 2d and Extended Data Fig. 9a,b). This interface is stabilized by a hydrophobic patch centered on β-NDC80 F74, β-NUF2 Y65, and α-NDC80 H110, together with adjacent acidic residues involving β-NDC80 E78 and β-NUF2 E80 that contribute electrostatic complementarity. Notably, NDC80 S69, an Aurora A phosphorylation site^59^, faces the negatively charged surface of the adjacent complex (Extended Data Fig. 9c). The interfacial residues are highly conserved across vertebrates but diverge in invertebrates (Extended Data Fig. 9d), consistent with budding yeast Ndc80C binding MTs with tubulin-dimer periodicity^20^, rather than the monomer-repeat arrangement.

Lowering the contour threshold revealed additional density extending from a previously uncharacterized kink (NDC80^288-292^/NUF2^170-174^) in the coiled-coil to an interhelical loop in a neighboring Ndc80C (Extended Data Fig. 10a). This kink is distinct from the canonical hinge region (NDC80^359-362^/NUF2^244-247^) and is absent in budding yeast^16,20,60-63^ (Extended Data Fig. 10b). AlphaFold3 (AF3)^64^ predicts a straight and continuous coiled-coil from the kink to the hinge (Extended Data Fig. 10a). Beyond the hinge, the distal Ndc80C coiled-coils (NDC80^363-642^/NUF2^248-464^), including the loop, are predicted by AF3 to form bundles with a spacing of ∼40 Å (Extended Data Fig. 10c), which matches both the tubulin repeat on MTs and the inter-Ndc80C spacing in our images (Extended Data Fig. 10d). Together, these features support a model in which a network of MT-stabilized contacts promotes cooperative Ndc80C clustering, thereby reinforcing robust KT-MT attachment.

### SkaC bridges tubulin interfaces on the MT lattice

Our cryo-EM map shows the SKA1 MTBD at a four-tubulin junction on the MT lattice, where it spans both the longitudinal interdimer interface along a PF, and the lateral interface between neighboring PFs (referred to as α1-, α2-, and β2-tubulins, Fig. 2e). The most extensive and well-defined contacts are formed with β2-tubulin, whereas the α1- and α2-tubulin interfaces are less well resolved (Fig. 2f-h and Extended Data Fig. 11a-d), suggesting local conformational flexibility. Overall, the SKA1-MT interface is dominated by basic SKA1 residues that contact acidic tubulin surfaces, with additional local hydrophobic reinforcement (Extended Data Fig. 11e,f). At the α1-tubulin interface, K183 and K184 lie opposite acidic residues in the α1-tubulin CTT, whereas the Aurora B site S185^27,29,33^ is positioned nearby within the same contact region (Fig. 2f and Supplementary Video 1). The adjacent α2-tubulin interface contains a positively charged SKA1 patch, including K223, K226, K227, and R155, and is further stabilized by a hydrophobic contact centered on SKA1 Y151-M152 and α2-tubulin Y262-P263 (Fig. 2g and Extended Data Fig. 11e). Notably, the Aurora B site T157 also maps close to this interface.^27,33^ At the β2-tubulin interface, basic residues R236, R239, and R245 contribute additional polar contacts near the αβ-tubulin interdimer junction, whereas P149, Y151, and M152 provide hydrophobic reinforcement (Fig. 2h and Extended Data Fig. 11f). The Aurora B site S242^27,33^ lies within this interface, whereas T205^27,33^ is positioned more peripherally, suggesting that phosphorylation at these sites could directly or allosterically modulate SkaC-MT affinity. These structurally defined contact sites provide an explanation for previous mutational analyses that identified the SKA1 basic patches and Aurora B phospho-sites within the MTBD as key determinants of MT binding and mitotic function^27,29,33,42^.

The SKA1 MTBD is separated from the hetero-trimerization and homo-dimerization region of the SkaC by a ∼40-residue unstructured segment (Extended Data Fig. 1a)^32,33^. Therefore, our reconstruction cannot define how the two MTBDs within a SkaC dimer are arranged relative to one another on the MT. When we mapped the SKA1 patches identified by cross-linking mass spectrometry (XL-MS) as being near tubulin onto our SKA1-MT structure, SKA1 K183/K184 are positioned >30 Å from β-tubulin, and K203/K206 lie >30 Å from α-tubulin^33^ (Extended Data Fig. 11g-i) and therefore do not contribute to the interaction of SKA1 with the MT (Fig. 2e-h). Instead, mutating these SKA1 residues was shown to abolish SKA1 binding to curved-tubulin^42^, suggesting that the SKA1 MTBD contains a distinct tubulin-interaction surface that recognizes curved tubulin.

### SkaC biases Ndc80C toward the extended MT lattice

Following polymerization, GTP hydrolysis induces conformational changes in tubulin that compact the MT lattice^65^, whereas growing MT ends contain GTP-bound tubulin and adopt a more extended conformation. The interaction of SKA1 across tubulin dimers at the longitudinal interdimer interface suggests that SkaC may be sensitive to the nucleotide-dependent state of the MT lattice (Fig. 2e). To test whether Ndc80C and SkaC exhibit distinct lattice preferences and whether their interaction alters these preferences, we used TIRF imaging to examine their binding to GMPCPP-capped GDP MTs, where GMPCPP-bound tubulin mimics the extended, GTP-like lattice at MT growing ends (Fig. 3)^65^.

**Fig. 3:**
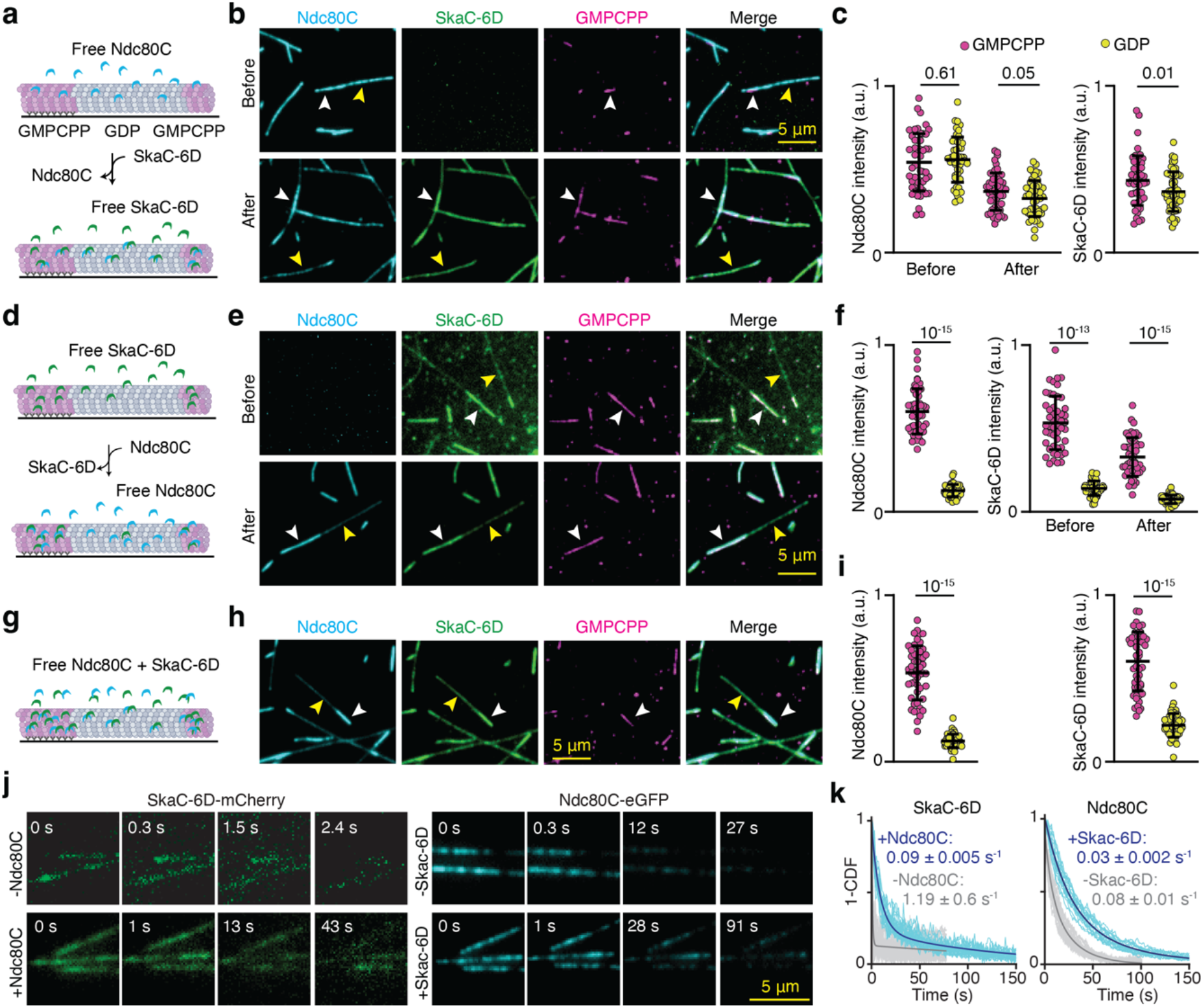
Ndc80C-SkaC exhibits lattice-selective binding to MTs. a, GMPCPP-capped GDP MTs were pre-incubated with Ndc80C before exchanging free Ndc80C in solution with SkaC-6D. **b-c**, MT binding (b) and fluorescence intensity (c) of Ndc80C and SkaC-6D on GDP (unlabeled) and GMPCPP (magenta) lattices before and after exchanging Ndc80C with SkaC-6D in solution. N = 50 MTs from two independent experiments. **d**, MTs were pre-incubated with SkaC-6D before exchanging free SkaC-6D in solution with Ndc80C. **e-f**, MT binding (e) and fluorescence intensity (f) of Ndc80C and SkaC-6D on GDP (unlabeled) and GMPCPP (magenta) lattices before and after exchanging SkaC-6D with Ndc80C in solution. N = 50 MTs from two independent experiments. **g**, Ndc80C and SkaC-6D were added simultaneously to MTs. **h-i**, MT binding (h) and fluorescence intensity (i) of Ndc80C and SkaC-6D on GDP (unlabeled) and GMPCPP (magenta) lattices when added simultaneously. N = 50 MTs from two independent experiments. **j**, Localization of SkaC-6D-mCherry and Ndc80C-eGFP on GMPCPP MTs after washing out unbound protein from the chamber (t = 0 s). **k**, The inverse cumulative distribution function (1-CDF) of SkaC-6D-mCherry and Ndc80C-eGFP intensity per length of an MT as time elapsed. Decay constants (±SE) were calculated from a fit to an exponential decay (solid curves, N = 20 MTs for each condition). In b, e, and h, white and yellow arrowheads indicate binding to the GMPCPP and GDP lattice, respectively. In c, f, and i, the center line and whiskers represent the mean ± SD. P-values are calculated from a two-tailed t-test.

Ndc80C alone bound similarly to GMPCPP and GDP lattices (Fig. 3a-c). Upon exchanging free Ndc80C in solution with SkaC-6D, the Ndc80C that remained bound on MTs recruited SkaC-6D to both GDP and GMPCPP lattices (Fig. 3a-c). In comparison, SkaC-6D alone exhibited weak binding to the GDP lattice but bound stably to the GMPCPP lattice (Fig. 3d-f), consistent with previous observations^42^. Upon introducing Ndc80C and removing free SkaC-6D, Ndc80C preferentially bound to the GMPCPP lattice through its interaction with lattice-bound SkaC-6D, while retaining the ability to bind independently to the GDP lattice, albeit with lower efficiency (Fig. 3d-f). When Ndc80C and SkaC-6D were introduced simultaneously, both proteins exhibited enhanced binding to the GMPCPP lattice compared with the GDP lattice (Fig. 3g-i). Together, these results indicate that SkaC preferentially recognizes the extended, GTP-like lattice and biases Ndc80C toward this lattice state, while Ndc80C promotes SkaC association with MTs independently of lattice state. This behavior is consistent with previous observations that Ndc80C promotes SkaC recruitment to kinetochores^23-25,27,34,35,66^ and that SkaC supports Ndc80C association with dynamic MT ends^29-31^.

Additionally, we measured dissociation kinetics of Ndc80C and SkaC from GMPCPP MTs after removing all unbound complexes from the flow chamber (Fig. 3j,k). The relatively rapid dissociation of SkaC in the absence of Ndc80C (Fig. 3j) was slowed by more than an order of magnitude in the presence of Ndc80C (Fig. 3j). Similarly, the presence of SkaC reduced the dissociation rate of Ndc80C by 3-fold (Fig. 3j). Together, these data support a reciprocal interaction in which SkaC biases Ndc80C towards the GTP-like lattice, while Ndc80C stabilizes SkaC association with MTs.

### SKA3 interacts with a vertebrate-conserved kink in the Ndc80C coiled-coil

Our structure shows the presence of a previously uncharacterized kink in the Ndc80C coiled-coil (NDC80^288-292^/NUF2^170-174^) that lies proximal to the canonical hinge (NDC80^359-362^/NUF^244-247^, Fig. 4a-d and Extended Data Fig. 10a,d). This kink is absent from the yeast Ndc80C (Extended Data Fig. 10b), and sequence conservation analysis suggests that it is more prominent in vertebrates (Fig. 4d). Interestingly, we observed additional density associated with this bending region, but it could not be directly assigned due to lower resolution (Fig. 4a). We integrated XL-MS constraints^30^ with iterative AF3 modelling of candidate SKA3 segments, which converged on a high-confidence interface between the Ndc80C kink region and SKA3 residues 355-380 (Fig. 4b and Extended Data Fig. 12a,b). The predicted SKA3-Ndc80C contact interface closely matched the cryo-EM density, despite differences in the bend and rotational angle of the Ndc80C coiled-coil region itself (Extended Data Fig. 12c). This assignment is consistent with prior evidence that neighboring regions within the SKA3 C-terminal domain participate in MT-associated interactions^30,67^ and that residues 355-380 contribute to Ndc80C binding in solution^31^. Our structure visualized its engagement with the Ndc80C kink region in the MT-bound state (Fig. 4a-c). The bound SKA3 segment includes a short α-helix (“tether” helix) and two functionally important phosphorylation sites, T358 and T360 (corresponding to D358 and D360 in SkaC-6D; Fig. 4c and Supplementary Video 1). The interface is organized around an extended hydrophobic zipper between SKA3 and NDC80/NUF2 (Fig. 4b), with several of the contributing hydrophobic residues highly conserved in metazoans (Fig. 4e). T358 (D358 in our structure) is located by NDC80 H320 and NUF2 K213 (Fig. 4c), the latter corresponding to a hotspot identified by XL-MS^30^.

**Fig. 4:**
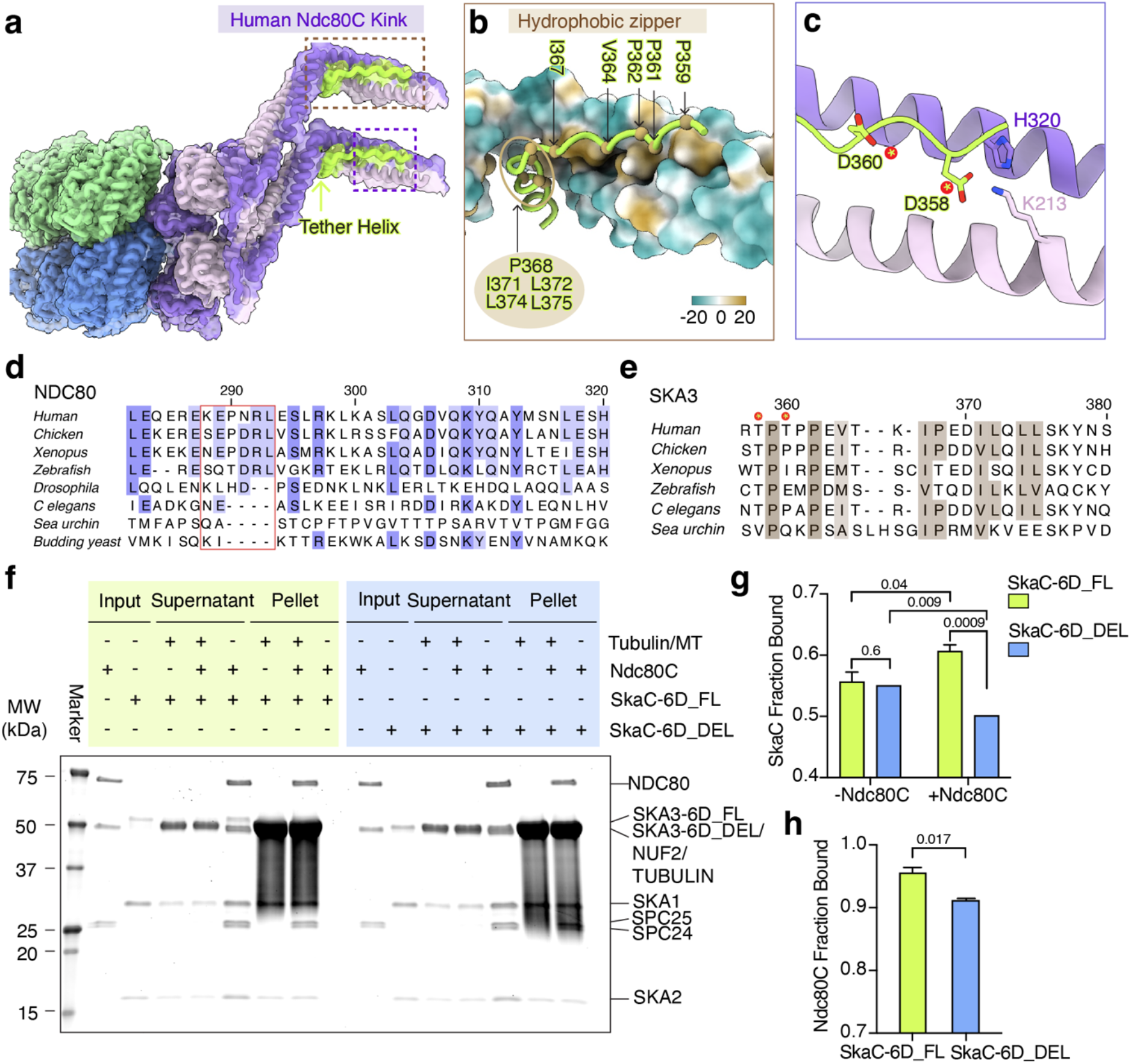
A phosphosite-containing SKA3 tether engages the Ndc80C kink. a, Cryo-EM density with fitted models showing the SKA3 tether helix and preceding region (residues 357-380) bound to the Ndc80C kink. Overview density is shown from the composite map, whereas the local SKA3-Ndc80C kink region density is shown from the Relion B-factor-sharpened map. **b**, Close-up view of the hydrophobic interface between the SKA3 segment and the Ndc80C kink. NDC80-NUF2 are shown as a hydrophobicity-colored surface, and SKA3 is shown as a ribbon with hydrophobic residues depicted as spheres. **c**, SKA3 residues T358 and T360 lie within the Ndc80C-binding interface and are represented here by the phosphomimetic substitutions D358 and D360 in the SkaC-6D construct used for EM studies. **d**, Sequence alignment of the Ndc80 kink and adjacent residues forming the SKA3-interacting interface shown in (**a**). Kink residues are boxed in red, and conserved interface residues are highlighted. Darker shading denotes higher sequence conservation. **e**, Sequence alignment of the SKA3 tether segment engaging the Ndc80 kink in (**a**) across the species in (**d**), excluding budding yeast. Hydrophobic residues at the interface positions are highlighted. **f**, Flamingo-stained SDS-PAGE analysis of MT-pelleting assays showing SkaC-6D_FL and SkaC-6D_DEL binding to MTs in the absence or presence of Ndc80C. **g,h**, Quantification of the MT binding ratios of SkaC (**g**) and Ndc80C (**h**) from the experiments shown in (**f)**. SkaC binding was quantified using the SKA2 band, and Ndc80C binding was quantified using the NDC80 band. Data are mean ± SD from three independent experiments, analyzed by a two-tailed t-test.

To validate this proposed interface, we generated a SkaC-6D mutant lacking this segment (SKA3 Δ355-380; SkaC-6D_DEL) and compared its MT-binding affinity with that of the full-length SkaC-6D (SkaC-6D_FL) in the presence or absence of Ndc80C (Fig. 4f-h). In MT pelleting assays, SkaC-6D_DEL and SkaC-6D_FL showed similar binding in the absence of Ndc80C (Fig. 4g), indicating that the deletion of this segment does not impair basal SkaC-MT association. By contrast, Ndc80C enhanced MT binding of SkaC-6D_FL, but reduced the binding of SkaC-6D_DEL, likely due to competition for an overlapping binding site on tubulin. Reciprocally, Ndc80C binding also diminished in the presence of SkaC-6D_DEL with respect to SkaC-6D_FL (Fig. 4h). These results indicate that the SKA3 355-380 segment is dispensable for basal SkaC-MT binding but is required for the cooperative binding of SkaC and Ndc80C on the MT surface. In the absence of this SkaC-Ndc80C coupling at the higher-radius coiled-coil region, the Ndc80C-CHD and SKA1-MTBD compete for MT binding, as indicated by the overlap on their tubulin footprint (Extended Data Fig. 5a,b and Supplementary Video 1). A concurrent study also identified the SKA3-binding interface at the Ndc80C coiled-coil kink^68^. Although the two studies differed slightly in SkaC preparation (6D versus CDK1-phosphorylated), MT stabilization (GMPCPP versus Taxol), the order of protein addition during grid preparation, and the assemblies retained for focused refinement of the kink region, their findings are highly consistent. Niu et al. showed that structure-guided mutations in HEC1/NDC80 and NUF2 disrupt SkaC binding^68^, in excellent agreement with our structural model andwith our complementary mutational analysis of the SKA3 tether, providing functional support from both sides (Ndc80C and SkaC) of the interface. Together, these findings identify the SKA3 355-380 segment as a tether that links SkaC to the Ndc80C coiled-coil kink, thereby enabling cooperative binding of the two complexes on MTs.

### SKA3 tethering couples Ndc80C-SkaC to MT dynamics

To determine whether the observed SKA3 tether-dependent differences in MT binding translate into changes in MT dynamics, we next examined the effects of Ndc80C and SkaC using TIRF microscopy (Fig. 5a,b). Compared to the control lacking either complex, both Ndc80C and SkaC individually stabilized MTs by increasing polymerization rate, reducing depolymerization rate, decreasing catastrophe frequency, and increasing rescue frequency (Fig. 5a,b). SkaC had a stronger effect at the concentrations used in our assay. In the presence of both complexes, SkaC-WT enhanced Ndc80C-mediated stabilization of dynamic MT ends. This effect was further amplified in the presence of SkaC-6D_FL (Fig. 5a,b), consistent with its phosphorylation-enhanced interaction with Ndc80C^31,34,39^.

**Fig. 5:**
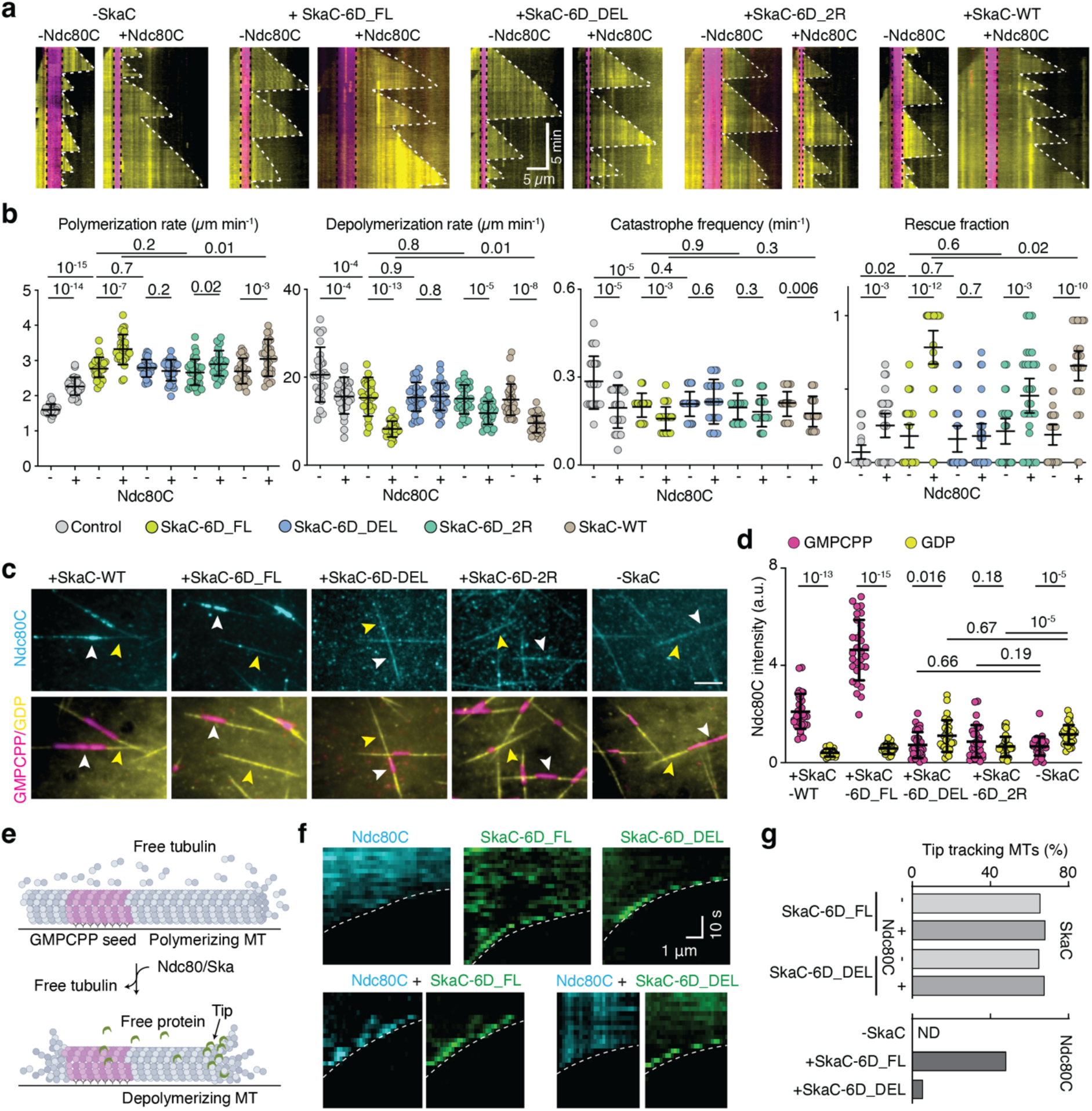
The SKA3 tether couples SkaC lattice selectivity to Ndc80C MT-end stabilization and tip tracking. a, Representative kymographs of dynamic MTs (yellow) in the presence of 50 nM Ndc80C-eGFP, 100 nM SkaC-6D or SkaC-6D mutants, 100 nM SkaC-WT alone or in combination, as indicated. Vertical black dashed lines mark the ends of GMPCPP-stabilized seeds (magenta), and white traces mark the MT plus ends. **b**, Quantification of MT plus-end polymerization rate, depolymerization rate, catastrophe frequency, and rescue fraction under the indicated conditions. The center line and whiskers represent mean ± SD for polymerization rate, depolymerization rate, catastrophe frequency and mean with 95% c.i. for rescue fraction. P values were calculated using a two-tailed t-test. N = 30 MTs were analyzed for each condition from two independent experiments. **c**, Representative images showing binding of Ndc80C-eGFP (50 nM) in the presence of Ska-WT and SkaC-6D (or its mutants) on GMPCPP MT lattices (magenta) versus GDP lattices (yellow). White and yellow arrowheads indicate binding on the GMPCPP and GDP lattices, respectively. Images were individually contrast-adjusted to visualize weak binding regions. Scale bar: 5 μm. **d**, Integrated fluorescence density of Ndc80C under the indicated conditions. The center line and whiskers represent mean ± SD, respectively. P values were calculated using a two-tailed t-test. N = 30 MTs were analyzed for each condition from two independent experiments. **e**, Schematic representation of the MT depolymerization assay. **f**, Representative kymographs of depolymerizing MTs showing Ska6D_FL and Ska6D_DEL (green) in the presence or absence of Ndc80C (cyan), as well as Ndc80C alone (cyan). MT plus ends are oriented to the right. **g**, Percentage of MTs exhibiting tip tracking by Ska6D_FL, Ska6D_DEL, and Ndc80C under the indicated conditions. ND: not determined. N = 35 MTs from two independent experiments.

To determine whether the observed SKA3 tether-dependent differences in MT binding affect MT dynamics, we tested a SKA3 deletion mutant lacking this tether segment (SkaC-6D_DEL). The absence of the tethering elements within SKA3 had no effect on MT stabilization compared to SkaC-6D_FL alone, but largely abolished the additional stabilizing effect observed together with Ndc80C (Fig. 5a,b). Mutation of the two phosphorylation sites within this region (T358R/T360R; SkaC-6D_2R) did not significantly alter the MT dynamics parameters observed for SkaC-6D_FL alone but exhibited an intermediate stabilizing effect relative to SkaC-6D_FL and SkaC-6D_DEL in the presence of Ndc80C (Fig. 5a,b). These results identify the SKA3 tether segment as a contributor to the cooperative suppression of MT plus-end dynamics by SkaC and Ndc80C, with the two phosphorylation sites within this region fine-tuning this activity.

### The SKA3 tether couples SkaC lattice selectivity to Ndc80C tip tracking

We next asked whether the SKA3 tether also transmits SkaC-dependent lattice-state recognition to Ndc80C. In the presence of SkaC-6D_FL, Ndc80C showed a stronger preference for the GTP-like lattice than in the presence of SkaC-WT (Fig. 5c,d), consistent with phosphorylation-enhanced SkaC-Ndc80C coupling^31,34,39^. By contrast, the lattice bias was abolished in the presence of SkaC-6D_DEL and substantially reduced in SkaC-6D_2R, indicating that deletion or mutation of the tether disrupts the coupling required to bias Ndc80C binding. These results indicate that the SKA1 MTBD contributes to the lattice-state preference of the Ska complex, while the SKA3 tether is important for coupling this preference to Ndc80C.

Because persistent coupling to dynamic MT ends is a defining feature of KT function^5,24,29,69^, we next tested whether the same tether provides the physical link through which SkaC confers plus-end tracking on Ndc80C^29^. SkaC-dependent tracking of Ndc80C-associated assemblies had been observed previously^29,31^, but the structural basis for this activity remains unknown. In our MT depolymerization assay, SkaC robustly tracked depolymerizing MT tips, whereas Ndc80C alone showed no detectable tip-tracking activity under the same conditions (Fig. 5e-g, and Supplementary Video 2). Full-length SkaC-6D (SkaC-6D_FL) recruited Ndc80C to depolymerizing MT ends and conferred tip-tracking behavior on Ndc80C. Deletion of the SKA3 tether (SkaC-6D_DEL) did not abolish SkaC tip tracking itself but largely prevented SkaC from conferring this activity on Ndc80C (Fig. 5e-g, and Supplementary Video 2). Together, these results identify the SKA3 tether as the structural conduit that transfers SkaC lattice selectivity and plus-end tracking to Ndc80C, converting Ndc80C into a Ska-coupled tip-tracking assembly.

## DISCUSSION

Our findings reveal how human SkaC couples Ndc80C to the structural state and dynamics of MT plus ends. In our structure, clustered Ndc80C on one PF positions SkaC on an adjacent PF through a phosphosite-containing SKA3 tether bound to a kink in the Ndc80C coiled coil. This arrangement brings the NDC80 CH domain and the SKA1 MTBD together around the α-CTT, forming a sandwich-like interface whose engagement is strengthened on polyglutamylated MTs. The SKA1 MTBD bridges the tubulin interdimer interface and preferentially recognizes the extended, GTP-like lattice characteristic of growing MT plus ends. Through the SKA3 tether, this SkaC-dependent lattice preference is transmitted to Ndc80C, promoting Ndc80C enrichment at the extended plus-end lattice, stabilization of dynamic MT ends, and Ndc80C tip tracking. Thus, the Ndc80C-SkaC interface provides a structural and regulatory mechanism for maintaining KT attachments that are strong enough to bear force yet adaptable enough to follow MT-end remodeling during chromosome segregation.

Our work identifies the tubulin tails as a tunable regulatory layer within the KT-MT attachment. Rather than serving only as flexible, negatively charged extensions of the MT surface, the α-tubulin C-terminal tail is structurally captured at the Ndc80C-SkaC interface, where its PTM state can tune assembly of the combined outer-KT MT-binding module. Removal of the tubulin tails weakens the binding of both Ndc80C and SkaC to MTs, and a stronger association to brain MTs versus HeLa MTs supports a positive contribution of tubulin PTMs to this interaction. Most specifically, the selective reduction of Ndc80C and SkaC recruitment on TTLL1-deficient, but not TTLL7-deficient MTs further argues that α-CTT polyglutamylation has a strongly positive effect in the binding of both Ndc80C and SkaC. These findings provide a molecular link between the tubulin code and outer-kinetochore attachment, suggesting that chromosome segregation can be regulated not only through phosphorylation of kinetochore proteins, but also through post-translational tuning of the microtubule lattice itself. Our data would also provide a mechanistic paradigm explaining previous findings demonstrating that loss of α-tubulin polyglutamylation catalyzed by TTLL11 leads to a high rate of chromosome segregation errors^44^, a common feature of kinetochore dysfunction^9,70^.

Our structures also clarify how human Ndc80C forms a cooperative MT-bound scaffold for SkaC engagement. Multivalent Ndc80C assemblies are required for efficient tip tracking and force coupling^69^, but the molecular basis for human Ndc80C cooperativity has remained incompletely defined. The NDC80 CHD engages the NDC80 NTT and NUF2 of a neighboring Ndc80C, templated by the MT lattice. Based on our results and previous studies^12,14,63,69^, we propose that the human Ndc80C cooperativity is nucleated via NDC80 CHD- and NTT-mediated interactions near the MT surface, and then propagates along the coiled-coil, generating a robust arrangement with the multivalency needed for sustained MT engagement and load bearing. This organization provides a metazoan solution to robust KT-MT coupling. Whereas budding-yeast Ndc80C gains multivalency through association with the oligomeric Dam1C ring^20^, human Ndc80C forms an intrinsic cooperative scaffold that positions dimeric SkaC on adjacent PFs. Thus, distinct eukaryotic lineages appear to have evolved different molecular architectures to meet the shared challenge of maintaining attachment to dynamic MT ends^1,2^.

Our structural findings provide a framework for understanding how mitotic phosphorylation regulates this attachment interface. Several regulatory phosphosites map directly to MT-binding, Ndc80C-Ndc80C, or Ndc80C-SkaC interfaces (Fig. 6a and Supplementary Video 1). NDC80 S165 defines a NEK2-regulated node at the CHD-MT interface, in line with its reported roles in chromosome alignment and spindle-checkpoint signaling^56-58^. Aurora B-sensitive sites in SKA1 within or immediately adjacent to its MT binding interface are consistent with Aurora B function in early-mitotic error correction and in restraining premature SkaC accumulation at KTs^27,29,33,71^. NDC80 S69 lies at the interface with a neighboring Ndc80C, providing a structural rationale for how Aurora A could dampen Ndc80 clustering to regulate metaphase KT-MT dynamics^59^, and consistent with MT-bound Ndc80 oligomerization to modulate Aurora kinase access to Ndc80C phosphosites^72^. Finally, SKA3 T358 and T360 map directly to the SKA3-Ndc80C kink interface, consistent with CDK1-dependent phosphorylation promoting SKA3 binding to Ndc80C and SkaC recruitment to KTs^31,34^. These observations suggest that mitotic kinases regulate KT-MT attachment not through a single switch, but by tuning multiple contact points within the Ndc80C-SkaC-MT assembly.

**Fig. 6:**
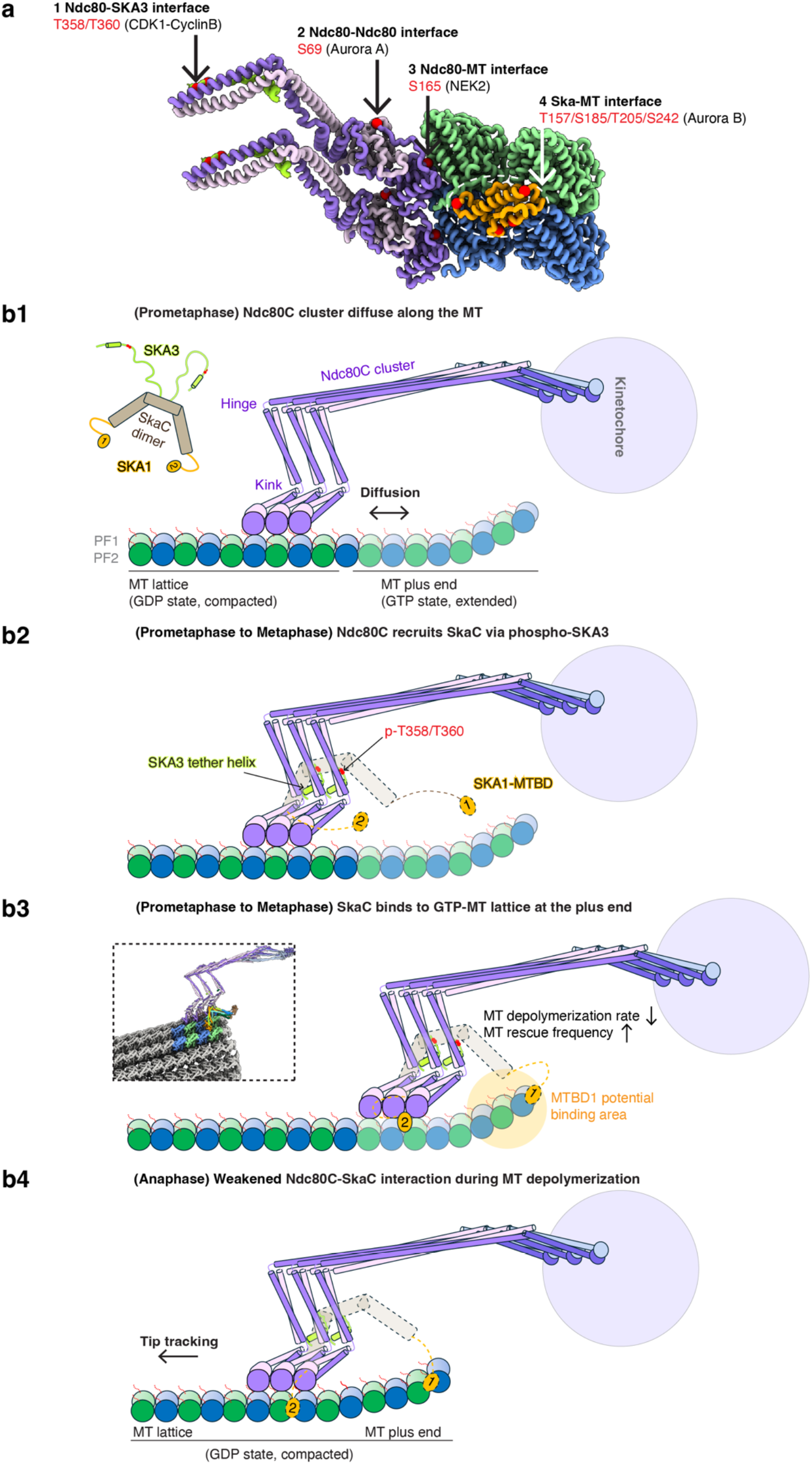
Regulatory phosphosites and a mechanistic model of Ndc80C-SkaC coordination at MT plus ends during mitosis. a, Structurally resolved regulatory phosphorylation sites (red spheres) relative to the MT-binding surfaces, adjacent Ndc80C interfaces, and the SKA3-Ndc80C interface (see also Supplementary Video 1). **b**, In prometaphase and metaphase, clustered Ndc80C recruits SkaC to adjacent PFs through the binding of the SKA3 tether segment to the Ndc80C kink region (**b1**-**b2**). SKA1 MTBD binds next to Ndc80C, stabilizes the MT end, and biases the assembly toward the extended GTP-like lattice at MT plus ends (**b3**). This coupling is expected to weaken in anaphase, as MT ends shorten and mitotic phosphorylation declines (**b4**). (α-tubulin (green), β-tubulin (blue), NDC80 (medium purple), NUF2 (thistle), SPC24 (slate blue), SPC25 (light blue)). SKA1 MTBDs are colored orange, and SKA3 C-terminal tails are colored green-yellow; SKA1-2-3 coiled-coil bundles are simplified as tan rectangles, and their positional uncertainty is indicated with dashed lines and reduced opacity. The dashed rectangle in B3 shows a more detailed model of the state described in our study.

The functional significance of this architecture is most apparent at dynamic MT plus ends, where KTs must remain attached while the underlying tubulin lattice continuously remodels. SkaC preferentially associates with the extended, GTP-like lattice by engaging the tubulin interdimer interface, whereas Ndc80C alone remains more broadly distributed on MTs. The observed lattice-state preference likely arises from hydrolysis-induced conformational changes in tubulin at polymerization interfaces^65,73^ that are recognized by the SKA1 MTBD. Other factors, such as differences in tubulin E-hook accessibility^74^, could also contribute to this preference. Through the SKA3 tether, this lattice preference is transmitted to Ndc80C, providing a mechanism for enriching the core force-coupling complex at the MT-end for persistent attachment. This interpretation helps explain previous observations that Ndc80C-bound SkaC stabilizes MT ends in a stalled state^31^, that SkaC can autonomously track growing plus ends^42^, and that a tripartite Ndc80-Cdt1-Ska complex supports processive bidirectional plus-end tracking^75^. Importantly, our tip-tracking assays revealed that the SKA3 tether provides a physical link that allows SkaC to transfer MT-end recognition to Ndc80C, enabling regulated attachment that can remain coupled to dynamic MT plus ends at KTs. Recent cellular work showed that perturbing an overlapping SKA3 C-terminal segment delays mitotic progression, supporting the physiological relevance of the SKA3 tether region structurally defined here^76^.

Previous cellular studies established that Ndc80C is recruited to KT early in mitosis before stable end-on attachments form^77,78^. In comparison, SkaC becomes strongly enriched following MT attachment^23,25,27^, increases progressively as KTs congress and biorient^26^, and reaches maximal levels at metaphase^27,79^. Our structural and functional data support a stepwise model for how coordinated Ndc80C-SkaC interactions maintain KT attachment to dynamic MT ends while modulating MT behavior (Fig. 6b). In prometaphase, KT-bound Ndc80C captures spindle MTs, forms cooperative clusters along PFs, and diffuses on the lattice (Fig. 6b1). As cells progress through prometaphase and metaphase, when CDK1 activity is high^80^, phosphorylated SkaC is recruited through the SKA3-Ndc80C kink interface (Fig. 6b2). In this arrangement, one SKA1 MTBD lies adjacent to Ndc80C and, together with the NDC80 CHD, stabilizes the α-CTT “sandwich” interface, and biases Ndc80C towards the extended GTP-like lattice at the MT plus end (Fig. 6b3). A second SKA1 MTBD may engage with the curved end-associated tubulin states, as previously proposed^33,42^, using a distinct surface from that used to interact with the straight MT lattice (Extended Data Fig. 11g-i). Through the SKA3 tether, this SkaC-dependent lattice preference is transmitted to Ndc80C, enabling Ndc80C tip tracking and stabilizing dynamic MT ends. This Ndc80C-SkaC arrangement is not necessarily a long-lived static assembly but can be dynamically renewed to maintain a coupling state at the MT interface^66,78^. As cells enter anaphase, depolymerizing MT ends and declining CDK1 activity^81^ may jointly disfavor SKA1 engagement with MTs and weaken phospho-dependent SKA3-Ndc80C interaction (Fig. 6b4), in line with the reduced KT abundance of SKA3 at this stage^82^. Together, this model integrates cooperative clustering, phospho-regulated recruitment, and lattice-state recognition for metazoan KTs to maintain persistent yet adaptable coupling to dynamic MT ends.

Our findings define the outer KT as a coordinated, multivalent molecular assembly that actively “reads” the lattice state to navigate the dynamic mitotic environment. Ultimately, this phospho-regulated and geometry-dependent coupling ensures the high-fidelity chromosome segregation necessary for organismal viability, while its disruption provides a mechanistic path toward the aneuploidy and chromosome instability observed in human disease^9,10,83^.

## DATA AND CODE AVAILABILITY

The motion-corrected cryo-EM micrographs have been deposited in EMPIAR under accession code EMPIAR-14018. The composite Tubulin-Ndc80-Ska map, the Tubulin-Ndc80-Ska consensus map, and focused maps of β-tubulin, two adjacent Ndc80 complexes, and the Ndc80 and SKA3 have been deposited in the Electron Microscopy Data Bank (EMDB) under accession codes 75942, 75938, 75939, 75940 and 75941, respectively. The coordinates for the corresponding models have been deposited in the Protein Data Bank (PDB) with accession code 11QA.

## ACKNOWLEDGMENTS

We thank Anthony Schuller for cloning of wild-type Ndc80 and Ska complex, Jie Fang for assistance with bacterial culture, Fabienne Pierre, Sinda Khanfir and Maria M. Magiera for technical support, Rui Zhang and Alfredo J. Florez for computational advice, Zhenlin Yang, Julia Peukes and Nikhit Kambdur for their assistance, Patricia Grob for EM support, Daniel Toso and Ravi Thakkar at the Cal-Cryo EM facility for help with EM data acquisition, Paul Tobias, Kurt Stine, and Victor Marquez for computing support, and Rui Zhang, Nicholas Lue and Juan Perez Bertoldi for their comments on this manuscript. The work was funded by NIH (GM127018 to EN and GM136414 to AY), NSF (MCB-2534447 to AY), the European Research Council (ERC) under the European Union’s Horizon 2020 research and innovation programme (Grant agreement ID: 101071583 ‘TubulinCode’ to EN and CJ), and EMBO Postdoctoral Fellowship (ALTF 1069-2024) to VS. EN is a Howard Hughes Medical Institute Investigator. We acknowledge the Imaging Methods Core Facility at BIOCEV, an institution supported by the MEYS CR (LM2023050 Czech-BioImaging), for their support and assistance in this work.

## AUTHOR CONTRIBUTIONS

J.Z and E.N. initially conceived the project; J.Z prepared protein samples, performed MT pelleting assays and EM studies, built and refined models, analyzed the structure; A.Y., I.S., Y.Z., E.N. and J.Z. designed TIRF experiments using porcine brain MTs; I.S. and Y. Z. performed TIRF experiments and analyzed data; C.J. and V.S. designed and performed TIRF experiments using MTs from mutant mouse brain and provided HeLa tubulin. E.N and A.Y. supervised the project and secured funding; J.Z. and E.N. wrote the initial manuscript, and E.N., J.Z., A.Y., I.S., C.J. and V.S. revised the manuscript with input from all authors.

## METHODS

### Cloning, expression, and purification of SkaC and Ndc80C

All SkaC and Ndc80C genes were cloned into the pETDuet vector, with or without an N-terminal 14×His-BD Sumo tag. SkaC mutants were generated via PCR-based site-directed mutagenesis or gene synthesis. All sequences were verified by whole plasmid sequencing prior to expression. Constructs were expressed in *E. coli* BL21(DE3) RIL or Rosetta2 (DE3) pLysS strains grown in Luria Broth supplemented with kanamycin, ampicillin, and chloramphenicol. Protein expression was induced with 0.4 mM isopropyl β-D-1-thiogalactopyranoside (IPTG) at 18 °C for 18-20 hours. Cells were harvested by centrifugation and resuspended in lysis buffer (25 mM Tris-HCl pH 8.0, 400 mM NaCl, 20 mM imidazole, 5% glycerol, 5 mM 2-mercaptoethanol) supplemented with protease inhibitor cocktail tablets and 1 mM PMSF. Cells were lysed by sonication and lysates were clarified by centrifugation at 40,000 rpm in a Ti45 rotor (Beckman) for 1 hour. The supernatant was incubated with Ni-NTA resin (GoldBio) for 2 hours, followed by washing with wash buffer (25 mM Tris-HCl pH 8.0, 150 mM NaCl, 40 mM imidazole, 5% glycerol, 5 mM 2-mercaptoethanol). The His tag was cleaved on-column overnight using Sumo protease. Proteins were further purified by ion-exchange chromatography using a HiTrap Q column (Cytiva), followed by size-exclusion chromatography on a Superdex 200 Increase 10/300 column (Cytiva) equilibrated with gel filtration buffer (25 mM Tris-HCl, pH 8.0, 150 mM NaCl, 5% glycerol, 1 mM DTT). Peak fractions were concentrated, flash frozen in liquid nitrogen, and stored at -80°C. The procedures above were used for all SkaC and Ndc80C purification.

### Tubulin purification

Tubulin was purified from wild-type and knock-out mouse brains or HeLa S3 cells grown in suspension using the polymerization/depolymerization protocol described previously^48^. In short, brains from wild-type (WT), Ttll1^-/-^, Ttll7^-/-^ or Ttll1^-/-^Ttll7^-/-^ mice were homogenized in lysis buffer (80 mM PIPES pH 6.9, 1 mM EGTA, 1 mM MgCl_2_, 1 mM 2-mercaptoethanol, 1 mM PMSF, and protease inhibitors), or HeLa cells were diluted in ice-cold lysis buffer and lysed using a French press. After homogenization, the lysates were ultracentrifuged (180,000 x g for brain tubulin or 112,000 x g for HeLa tubulin at 4 °C for 30 min), the supernatant was used for the first polymerization round in 1 mM GTP, 30% glycerol at 30 °C for 30 min. Polymerized MTs were pelleted at 180,000 x g, 30 °C for 30 min and subsequently depolymerized on ice for 20 min. The second polymerization/depolymerization cycle was performed in 0.5 M PIPES-KOH pH 6.8 buffer and the third cycle was performed as the first cycle. Purified tubulin was snap frozen and kept at -80 °C until MT assembly.

### MT polymerization

MT preparation for EM sample: Porcine brain tubulin (Cytoskeleton) was reconstituted to 10 mg/ml in BRB80 buffer (80 mM PIPES, pH 6.9, 1 mM ethylene glycol tetraacetic acid (EGTA), pH 8.0, 1 mM MgCl₂) supplemented with 10% (v/v) glycerol, 1 mM GTP, and 1 mM DTT. Tubulin polymerization was initiated by incubating 10 µL of the solution at 37 °C for 30 minutes. MTs were pelleted by centrifugation at 20,000 × *g* for 20 minutes. The supernatant was carefully removed, and the pellet was resuspended in 8 µL of cold BRB80 buffer supplemented with 1.25 mM GMPCPP (Jena Bioscience). The sample was incubated on ice for 60 minutes to facilitate complete GTP-to-GMPCPP exchange. GMPCPP-stabilized MTs were then repolymerized at 37°C for 30 minutes. 8 µL of the polymerized sample was mixed with 52 µL of warm BRB80 buffer and centrifuged at 20,000 × *g* for 20 min at 37 °C. The supernatant was removed, and the pellet was resuspended in 45 µL of warm BRB80 buffer. The tubulin concentration was determined by depolymerizing MTs using 0.1 M CaCl₂ on ice for 10 min. Finally, GMPCPP-stabilized MTs were aliquoted into 5 µL portions, flash-frozen in liquid nitrogen, and stored at -80 °C.

MT preparation for pelleting assays: Tubulin aliquots were thawed and centrifuged at 10,000 × g for 10 min to remove aggregates, and the supernatant was collected for further use. GMPCPP-stabilized MTs were polymerized in cold BRB80 containing 40 µM tubulin, 1.25 mM GMPCPP, 10% glycerol, 1 mM MgCl₂ and 1 mM DTT, followed by incubation at 37 °C for 1 h. Polymerized MTs were pelleted at 10,000g for 15 min, resuspended in prewarmed BRB80, and kept at room temperature until use.

MT preparation for TIRF assay with mutant mouse tubulin: GMPCPP-stabilized MTs were polymerized from desired tubulin (WT or PTM variant tubulin) by incubation 4 mg/ml of tubulin for 30 min at 37 °C in BRB80 (80mM PIPES, 1mM EGTA, 1mM MgCl_2_, pH 6.9) supplemented with 1 mM MgCl_2_ and 1 mM GMPCPP (Jena bioscience). Polymerized MTs were diluted in BRB80 and pelleted at 18000 x g in a Microfuge 18 Centrifuge (Beckman Coulter). After centrifugation, MTs were resuspended in fresh BRB80 and kept at room temperature until usage.

### MT subtilisin treatment

Subtilisin-treated MTs were prepared by incubating 20 µM GMPCPP-stabilized MTs with 100 µg ml⁻¹ subtilisin (Sigma-Aldrich) at 37 °C for 10 or 120 min. Reactions were terminated by addition of 2 mM PMSF. Cleaved MTs were then pelleted at 10,000 × g for 10 min at 25 °C and resuspended in 20 µL prewarmed BRB80 buffer.

### MT pelleting assay and statistics analysis

Purified SkaC and Ndc80C were buffer-exchanged into BRB80 supplemented with 50 mM KCl and 10% glycerol using Micro Spin Desalting Columns (ThermoFisher). Pelleting assays were performed using 2 µM MTs mixed with 200 nM SkaC, with or without 200 nM Ndc80C, in 20 µL reaction volumes. After a 10 min incubation at room temperature, samples were centrifuged at 100,000 × g for 10 min at 25 °C. 15 µL of the supernatant was removed from the side opposite the pellet, mixed with 4× Laemmli sample buffer (Bio-Rad) and boiled for 5 min. Pellets were washed once by gently adding 50 µL BRB80, discarding the wash, and resuspending the pellet in 20 µL ice-cold BRB80 containing 50 mM CaCl_2_. After 10 min on ice, 15 µL of the suspension was mixed with 4× sample buffer and boiled for 5 min, followed by SDS-PAGE analysis.

Gels were fixed in 40% (v/v) methanol and 10% (v/v) acetic acid for 2 h, stained with Flamingo fluorescent dye (Bio-Rad) for 3 h, briefly rinsed in 0.1 % (w/v) Tween-20 for 10 min to reduce background and imaged on a ChemiDoc MP system (Bio-Rad). Band intensities were quantified using Image Lab (Bio-Rad). MT-bound proteins co-sediment with MTs and appear in the pellet, while unbound proteins stay in the supernatant. SkaC binding was quantified using the SKA2 band, and Ndc80C binding was quantified using the NDC80 band, as these bands were well resolved from the other subunits and allowed reliable densitometric measurement. The bound fraction was calculated from densitometric intensities as pellet / (pellet + supernatant). Under our assay conditions, SkaC or Ndc80C alone did not sediment in the absence of MTs. Data are mean ± SD from three independent experiments. Statistical significance was assessed with two-tailed t-test.

Because HeLa and porcine brain tubulin showed different polymerization efficiencies under identical assembly conditions, with HeLa tubulin consistently yielding a higher tubulin pellet fraction than porcine brain tubulin. To account for differences in the amount of assembled MTs present in each reaction, SkaC binding was normalized to the MT pellet fraction. Specifically, the fraction of SkaC recovered in the pellet was calculated as SkaC_pellet_ / (SkaC_pellet_ + SkaC_supernatant_), and this value was divided by the fraction of tubulin recovered in the pellet, Tubulin_pellet_ / (Tubulin_pellet_ + Tubulin_supernatant_), to obtain the MT-normalized SkaC fraction bound. MT-normalized Ndc80C fraction bound was calculated in the same way. Data are mean ± SD from three independent experiments. Statistical significance was assessed with two-tailed t-test.

### Cryo-EM grid preparation and data acquisition

To prepare SkaC-Ndc80C-MT samples for cryo-EM analysis, a frozen aliquot of GMPCPP-stabilized MTs was thawed at 37°C for 15 minutes and diluted to a final concentration of 2.5 µM. A 4 µL aliquot of MTs was applied to a glow-discharged holey carbon grid (QuantiFoil Au R 1.2/1.3, 200 or 300 mesh) and incubated for 1 minute. After manual blotting with Whatman #1 filter paper, 3 µL of SkaC was added, followed by 1 µL of Ndc80C, which was incubated for 1 minute. The grid was then transferred to a Vitrobot (ThermoFisher) set at 25°C and 100% humidity. Blotting was performed with a force of 5 pN and a blot time of 7 seconds before plunge-freezing in liquid ethane. Samples were stored in liquid nitrogen for later use. Cryo-EM data were collected using a Titan Krios G3i microscope (ThermoFisher) operating at 300 kV and equipped with a BIO Quantum Energy Filter (Gatan). Images were recorded on a K3 direct electron detector (Gatan) at a nominal magnification of 81,000×, corresponding to a calibrated physical pixel size of 1.048 Å. Data were acquired in super-resolution mode with a dose rate of 8.003 e⁻/px/s on the detector. Each exposure lasted ∼7.4 seconds and was dose-fractionated into 50 frames, corresponding to a total dose of ∼50 e⁻/Å² on the specimen. Data collection was performed semi-automatically using the SerialEM software suite^84^ and parameters are summarized in Table S1.

Assembly of MT-Ndc80C-only was performed under identical conditions, except that the 2ul of Ndc80C added directly to MT, without SkaC. Cryo-EM data were collected using an Arctica microscope (ThermoFisher) operating at 200 kV. Images were recorded on a K3 direct electron detector (Gatan) at a nominal magnification of 36,000×, corresponding to a calibrated physical pixel size of 1.14 Å. The camera was operated in super-resolution mode at a dose rate of ∼7.2 electrons/pixel/s on the detector. An exposure time of ∼9 s was used, dose-fractionated into 50 frames, corresponding to a total accumulated dose of ∼50 electrons/Å² on the specimen. Data collection was performed semi-automatically using the SerialEM software suite^84^.

### Cryo-EM data processing

MT-Ndc80C-SkaC data processing: A total of 17,066 movie stacks were motion-corrected and dose-weighted using the RELION 3.0’s built-in implementation^85,86^. The contrast transfer function (CTF) parameters for each micrograph were estimated using Ctffind4^87^. As previously described^88,89^. MTs were manually picked in RELION 3.0. MT segments were extracted along the length of each MT using a box size of 512 pixels, with neighboring boxes spaced 82 Å apart. The PF number of each MT was determined through supervised 3D classification, using low-pass-filtered (20 Å) references of 13- and 14-pf MTs as initial models. A total of 92% of GMPCPP-MTs were classified as 14-pf MTs and selected for further processing. Global alignment parameters were determined in RELION 3.0 using a low-pass-filtered 14-pf MT map as the initial reference. The resulting alignment parameters were then imported into FREALIGN for seam refinement and further processing^90^, following our previously published protocols^91^. Refined alignment parameters from FREALIGN were subsequently imported into RELION 3.0 for symmetry expansion, after which all further processing steps were performed using RELION 5.0.

A sub-region covering 3 longitudinal and 4 lateral tubulin dimers was re-extracted with a smaller box size of 256 pixels,^92^ yielding 5,872,504 particles after symmetry expansion. Signal-subtracted datasets targeting (i) a single dimer, (ii) two longitudinal dimers, or (iii) two lateral dimers were then subjected to focused 3D classification to resolve Ndc80C-SkaC binding modes:

(i) Single dimer: Three major classes were identified: Ndc80C bound to the tubulin monomer repeat, SkaC bound to the tubulin interdimer repeat, and a poorly bound or unbound class. The Ndc80C-bound class, comprising 1,842,582 particles, was selected as the starting point for Part III processing.
(ii) Two longitudinal dimers: The classification yielded three major classes, similar to those observed for the single dimer.
(iii) Two lateral dimers: Focused 3D classification revealed three major classes: Ndc80C bound to two lateral tubulin dimers, two poorly or non-decorated lateral tubulin dimers, and SkaC and Ndc80C co-bound to two lateral tubulin dimers. The SkaC and Ndc80C co-bound class, containing 957,949 particles, was used as the starting point for Part I processing (see Part I processing details below).

Part I Tubulin-Ndc80C-SkaC complex: A total of 957,949 particles from the SkaC and Ndc80C co-bound class were selected after two rounds of lateral tubulin dimer classification. These particles were re-extracted and subjected to one round of local refinement. Next, we generated a soft mask encompassing the two lateral tubulin dimers and their associated Ndc80C and SkaC densities, which was applied for particle signal subtraction. The subtracted particles were then subjected to focused 3D classification without alignment, with Blush Regularization enabled (K = 4). A small class comprising 17.9% of particles, containing only SkaC-bound complexes, was removed. The remaining particles were reverted to their original coordinates and subjected to another round of local refinement. A second round of focused 3D classification without alignment was then performed using smaller subtracted particles, and a minor class (12.4%) containing only Ndc80C-bound complexes was excluded. This resulted in a clean dataset of SkaC and Ndc80C co-bound particles. Using the same strategy, a few more cycles of local refinement and focused 3D classification with progressively smaller masks were carried out, leading to improved visualization of structural features, such as the α-CTT. Final local refinement including 154,557 particles yielded a 2.8 Å density map of the SkaC-Ndc80C co-bound state. The final cryo-EM map was sharpened using DeepEMhancer^93^.

Part II β-CTT: To improve the density of the β-CTT, we used the pool of 957,949 particles from the SkaC and Ndc80C co-bound class (used as a starting point for Part I). Focused 3D classification was performed using subtracted particles containing a single tubulin dimer, and 29.7% of particles with poor or lacking density for Ndc80C were removed. The remaining particles were reverted to their original coordinates and subjected to one round of local refinement. Next, a mask covering only the β-CTT tail and its surrounding region was generated, and particle subtraction was performed using this mask. Another round of focused 3D classification with subtracted particles was carried out, and a class representing 49.2% of particles with clear tail density was selected and reverted. After an additional local refinement with these reverted particles, a final round of focused 3D classification around the β-CTT region yielded a class with well-resolved β-CTT density. A total of 176,590 particles were selected for the final reconstruction, resulting in a map with an overall resolution of 3.4 Å. The final cryo-EM map was post-processed using DeepEMhancer for sharpening.

Part III Two adjacent Ndc80C coiled coils: To improve density in the coiled-coil and kink regions, we began with 1,842,582 particles from the one-tubulin-dimer classification. Two adjacent Ndc80C densities (without tubulin signal) were subtracted, followed by one round of focused 3D classification on the subtracted particles. The best class showing clear density for both α- and β-Ndc80C was selected, reverted to original coordinates, and subjected to one round of local refinement. We then re-centered and re-extracted these particles, followed by another round of local refinement. Next, two adjacent Ndc80C densities were subtracted again, and a further round of focused 3D classification was performed. A class comprising 37.8% of particles with improved coiled-coil density was selected, and 194,299 subtracted particles were refined to yield a density map containing two adjacent Ndc80 molecules at an overall resolution of 3.3 Å. The resulting map was sharpened in DeepEMhancer.

Part IV Ndc80C kink (with SKA3): For better density at the kink, we continued with the 194,299 subtracted particles from the last step of Part III. Particle subtraction was performed separately on the α- and β-Ndc80C, then combined, yielding 388,598 single Ndc80C particles for local refinement. A mask was created around the kink region, and focused 3D classification was carried out (K=3, T=80). A class comprising 34.0% of particles with a distinct helix under the kink was identified; these 115,893 particles were selected for a final round of local refinement. The resulting map was sharpened in RELION 5.0.

Part V α2-CTT: We started with 957,949 particles from the Ndc80C and SkaC co-bound class (same set used as a starting point in Parts I and II). These particles were re-extracted and recentered on the α2-tubulin tail, followed by one round of local refinement. Next, particle subtraction was performed using a mask covering the two lateral tubulin dimers, and particles from two classes displaying Ndc80C density on the right PF were selected and reverted. These reverted particles were then subjected to another round of local refinement. We then performed an additional round of particle subtraction and focused 3D classification, where 37.8% of particles displayed clear density extending from the α2-CTT to the Ndc80C on the right PF. After reversion, a reconstruction map was obtained at an overall resolution of 3.0 Å using 247,515 particles.

MT-Ndc80C-only data processing: 990 movie stacks were collected and processed using the same procedure for motion correction and CTF estimation. MT segments were extracted along the length of each filament after manual picking, using a box size of 1024 pixels with adjacent boxes spaced 82 Å apart. A total of 32,866 particles were then subjected to 2D classification in RELION 3.0.

### Cryo-EM model building, refinement and validation

Tubulin dimers from a previously published structure (PDB: 6DPU)^94^ were rigid-body fitted into the cryo-EM density map using UCSF Chimera or Chimera X^95,96^. Because the C-terminal tail regions are less well-resolved and contain residues that vary among tubulin isoforms, they were modeled as alanine residues, except for a few positions that are 100% conserved. For the SkaC, a complete dimeric structure was predicted using AF3^64^, and the MTBD from this prediction was manually placed within the density as an initial fitting step. For the Ndc80C, relevant fragments were similarly generated by AF3 and docked into the density map.

Following these initial placements, all models were manually adjusted in Coot to correct local discrepancies with the density^97^, and further interactive model refinement was performed using ISOLDE^98^. Final real-space refinement was carried out in Phenix to optimize model geometry and ensure optimal fit to the experimental map^99^. Maps were sharpened using postprocessing in RELION and DeepEMhancer. Local, sharpened maps were then fitted to the consensus map and merged into a single composite map of Tubulin-Ndc80-Ska using the “vop maximum” command in ChimeraX. Both the RELION and DeepEMhancer maps were used for model building, whereas only the RELION map was used for model refinement. All validation and refinement statistics are summarized in Supplementary Table 1.

### Preparation of GMPCPP-stabilized MT seeds

MT seeds were stabilized with a slowly hydrolysable GTP analogue, GMP-CPP. The ultracentrifuge, rotor, tubes, and BRB80 buffer (80 mM PIPES, 1 mM MgCl_2_, 1 mM EGTA, 1 mM DTT, pH 6.8 adjusted with KOH) were prechilled. Tubulin was mixed with 5% biotin-tubulin and 7% Cy5-labeled tubulin, diluted to 15 mg mL^-1^ in cold BRB80, and incubated on ice for 10 min. The mixture was cold spun at 400,000 *g* for 10 min to remove inactive tubulin. The supernatant was transferred to a fresh tube and GMP-CPP was added to a final concentration of 1mM followed by addition of DMSO to a final concentration of 10% and incubated at 37 °C for 30 min. After equilibrating the centrifuge and buffer to 37 °C, the sample was spun again at 400,000 *g* for 10 min. The pellet was gently resuspended in 30 µL of warm BRB80 using a cut pipette tip to prevent shearing. Aliquots of GMPCPP-stabilized seeds were snap-frozen and stored at -80 °C.

### GMPCPP-capped GDP MT Assays

Tubulin was clarified by ultracentrifugation at 400,000 × g for 10 min at 4 °C. GMPCPP-stabilized MT seeds were polymerized using a mixture of unlabeled, biotin-labeled, and Cy5-labeled tubulin and immobilized on biotin-PEG-coated coverslips through streptavidin linkage. GDP MT extensions were polymerized from the GMPCPP seeds using 1.7 mg/mL tubulin (containing 7% Cy3-labeled tubulin) in polymerization buffer (1 mg/mL casein, 0.5% Pluronic F-127, 50 mM KCl, 1 mM DTT, 1 mM GTP, and 0.2% methylcellulose in BRB80) for 15 min on a 37 °C heating stage (Oko Labs). GDP MTs were then capped by extending their ends for 30 s with 0.3 mg/mL unlabeled tubulin in capping buffer (1 mg/mL casein, 0.5% Pluronic F-127, 50 mM KCl, 1 mM DTT, 1 mM GMPCPP, and 0.2% methylcellulose in BRB80). The chamber was subsequently washed with wash buffer (1 mg/mL casein, 0.5% Pluronic F-127, 50 mM KCl, 1 mM DTT, and 0.2% methylcellulose in BRB80) to remove free tubulin and allow uncapped GDP MT extensions to depolymerize back to GMPCPP seeds.

MTs were pre-decorated with 300 nM SkaC-6D-mCherry or 150 nM Ndc80C-eGFP in imaging buffer (1 mg/mL casein, 0.5% Pluronic F-127, 1 mM DTT, 0.2% methylcellulose, 0.1 mg/mL glucose oxidase, 0.2 mg/mL catalase, and 0.8% D-glucose in BRB80). During imaging, the solution containing the first protein was exchanged with imaging buffer containing the second protein. In separate experiments, SkaC-6D-mCherry and Ndc80C-eGFP were introduced simultaneously into the chamber. Static images were acquired at multiple positions within each chamber, and fluorescence intensities along the MTs were quantified using Fiji.

### Single-molecule dissociation kinetics

Flow chambers were assembled by sandwiching a glass slide (drilled with two holes) and a PEG/PEG-biotin-coated coverslip using permanent double-sided tape. GMPCPP-stabilized MTs were immobilized on the biotin-PEG-coated surface via streptavidin linkage. Imaging chambers were then filled imaging buffer (1 mg mL-1 casein, 0.5% Pluronic F-127, 0.5 mM DTT, 0.2% methylcellulose, 0.1 mg/mL glucose oxidase, 0.2 mg/mL catalase, 0.8% D-glucose in BRB80 pH 6.8) containing SkaC-6D-mCherry (180 nM when alone or 100 nM when in presence of Ndc80C-eGFP) and/or 50nM Ndc80C-eGFP. After acquisition of ∼15 frames, the chamber was washed with imaging buffer during continued image acquisition to monitor the bulk detachment of the proteins from the GMPCPP MTs.

### Dynamic microtubule imaging

Tubulin was clarified by ultracentrifugation at 400,000 g for 10 min at 4 °C. Biotinylated GMPCPP-stabilized seeds were immobilized on biotin-PEG-coated coverslips as described above. Imaging chambers were filled with a dynamic MT mixture (1 mg/mL casein, 0.5% Pluronic F-127, 0.5 mM DTT, 2 mM GTP, 0.2% methylcellulose, 0.1 mg/mL glucose oxidase, 0.2 mg/mL catalase, 0.8% D-glucose in BRB80 pH 6.8) containing 1.5 mg/mL tubulin (contains 7% of Cy3-labelled tubulin) and 100 nM SkaC (or its mutants) and/or 50 nM Ndc80C-eGFP. The chamber was then sealed with nail polish to prevent evaporation during imaging. The assay was performed at 33 °C. Images were captured every 5 s for 150 frames per imaging area and analyzed as kymographs made in FIJI. For MT lattice-selection assays with Ndc80C-eGFP and SkaC-6D-mCherry, unlabeled tubulin was used in all conditions except SkaC-6D-mCherry alone, for which 7% Alexa488-labeled tubulin was included.

To test tip tracking of SkaC and Ndc80C, MT depolymerization was triggered by removal of soluble tubulin with imaging buffer (1 mg/mL casein, 0.5% Pluronic F-127, 0.5 mM DTT, 0.2% methylcellulose, 0.1 mg/mL glucose oxidase, 0.2 mg/mL catalase, 0.8% D-glucose in BRB80 pH 6.8). Assays were performed in the presence of 25 nM LD655-labeled SkaC-6D_FL-SNAP and SkaC-6D_DEL-SNAP and 30 nM Ndc80C-eGFP. Images were captured every 200 ms and analyzed as kymographs in FIJI.

### TIRF Microscopy

For the dynamic MT assay, total internal reflection fluorescence (TIRF) microscopy was performed on a custom-built multicolor objective-type TIRF microscope based on a Nikon Ti-E body equipped with a 100×, 1.49 N.A. oil-immersion apochromatic objective and Perfect Focus System. Fluorescence was detected using an electron-multiplying CCD camera (Ixon EM+, Andor; 512 × 512 pixels) with an effective pixel size of 160 nm after magnification. eGFP, Cy3 or mCherry, and LD655 fluorophores were excited with 488, 561, and 633 nm lasers (Coherent) coupled through a single-mode fiber (Oz Optics), and emission was collected through 525/50, 593/46 (Cy3) or 629/56 (mCherry), and 697/75 nm bandpass filters (Semrock), respectively. The system was controlled with Micro-Manager v1.4. For the TIRF assays using mutant mouse tubulin, TIRF microscopy assays were performed on an inverted widefield microscope (Nikon Eclipse Ti2), equipped with a motorized XY-stage, transmitted light lamp (Nikon TI2-D-LHLED), Nikon Apo TIRF 60x oil immersion objective (NA 1.49), 1.5x tube lens, and PRIME BSI camera (Teledyne Photometrics). MTs were visualized using interference reflection microscopy (IRM), and Ndc80-eGFP and Ska-mCherry were imaged using EM435-485 and EM500-545 filter turrets, respectively.

### TIRF microscopy with mouse mutant and wild-type brain tubulin

TIRF chambers were assembled by melting thin parafilm strips in between two glass coverslips silanized with hexamethyldisilazane (HMDS, Sigma, #379212). Chambers were incubated with 20 µg/mL anti-β-tubulin antibodies (#T7816, Sigma) and passivated with 1% F-127 (#P2443, Sigma, 1% in PBS). The PTM-variant MTs were immobilized into the chamber, identified using IRM, after which WT MTs were immobilized. Before imaging, chambers were equilibrated in TIRF assay buffer AB (BRB80 pH 6.9, supplemented with 10 mM dithiothreitol, 0.02 mg/ml casein, 1 mM Mg-ATP, 20 mM D-glucose, 0.1% Tween20, 0.22 mg/ml glucose oxidase and 20 µg/ml catalase). Next, Ndc80C-eGFP and SkaC-mCherry were diluted to 4 nM in AB, added to the chamber, and imaged using TIRF microscopy.

### Data analysis for TIRF microscopy

Dynamic MTs were captured for 150 frames per imaging area and analyzed as kymographs in FIJI. Background-subtracted integrated fluorescence intensity of Ndc80C-eGFP and SkaC-6D-mCherry on the GMPCPP or GDP MT was calculated in FIJI. Catastrophe frequency was defined as the number of plus-end catastrophe events on each MT divided by the elapsed time. Rescue fraction was defined as the number of plus-end rescue events on each MT divided by the total number of catastrophes on the same MT.

For the off-rate measurement experiment, the integrated fluorescence intensity was calculated from each MT and corrected for photobleaching using a custom-written macro in FIJI. The change in integrated fluorescence intensity over time was fitted to a bi-exponential decay function in MATLAB. The rate constants were defined as the weighted average of the two decay rates obtained from the fit. P values were calculated from the two-tailed Student’s t-test in GraphPad Prism.

In the mouse mutant tubulin and concentration dependency assay, Ndc80C-eGFP or SkaC-mCherry density on the different MT variants was measured after 5 minutes of incubation and quantified using ImageJ2 (2.16.0/1.54g). For each experiment, Ndc80C or SkaC density was measured as the mean intensity of a 5-pixel-wide line drawn along the microtubule in the 488- or 561-channel, respectively. Background was estimated for each MT by drawing an identical line directly adjacent to the MT, and the background mean was subtracted from the MT mean. For the mouse mutant tubulin assay, the Ndc80C or SkaC density on the PTM-variant MTs was then normalized to the average density on the WT MTs of the same experiment.

**Extended Data Fig. 1:**
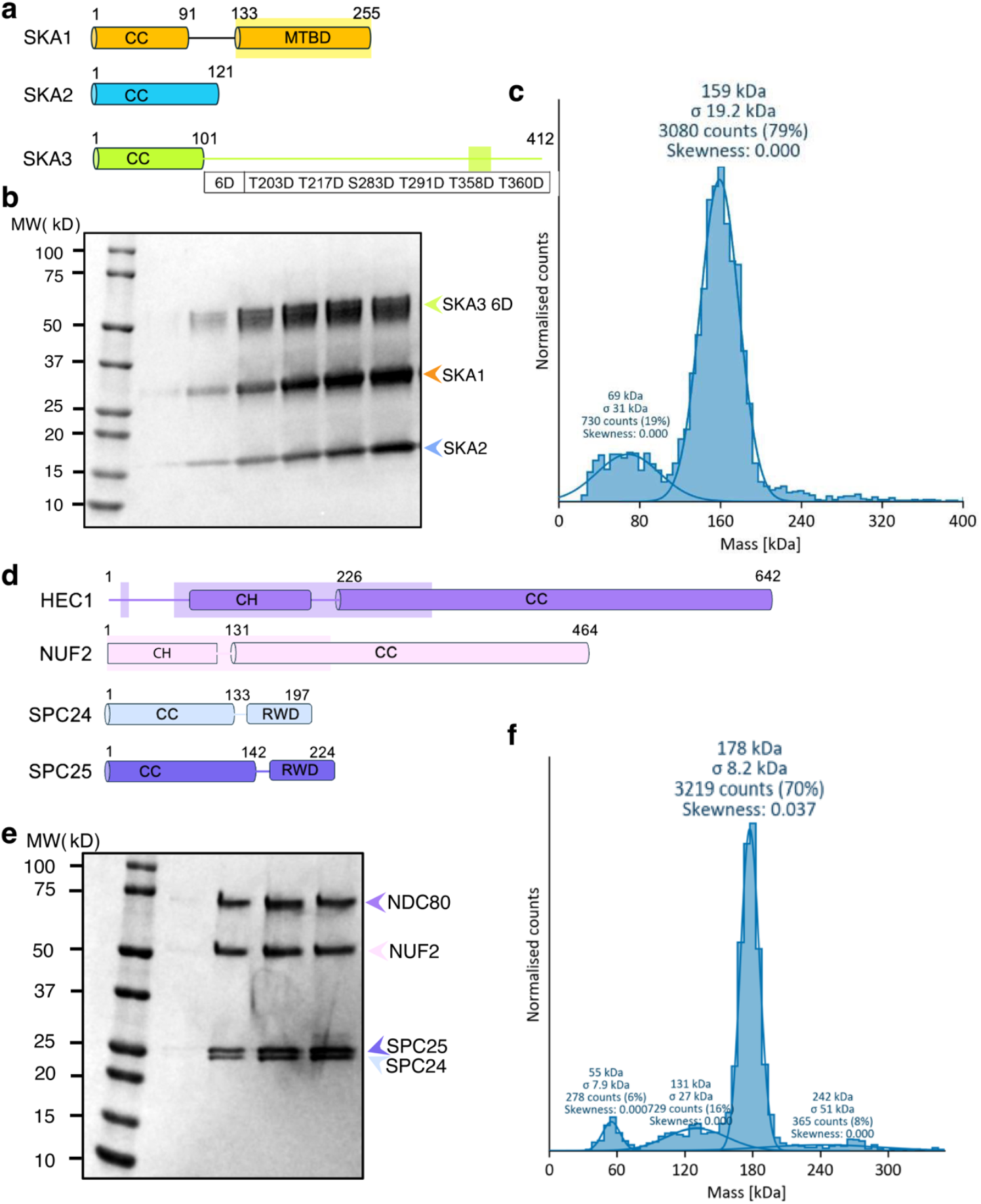
Purification and characterization of the SkaC-6D and Ndc80C samples, used for cryo-EM analyses. a, Schematic of the human SkaC-6D construct, with phosphomimetic substitutions in SKA3 at T203, T217, S283, T291, T358, and T360. Regions resolved in the cryo-EM maps are shaded (SKA1: 136-255; SKA3: 357-380). **b**, SDS-PAGE gel of SkaC-6D after gel filtration. Bands corresponding to the three SkaC subunits are indicated. **c**, Mass photometry analysis of SkaC-6D shows a major peak at the expected molecular mass for the SkaC dimeric complex. The measured molecular masses, associated errors, and relative populations are indicated. **d**, Schematic of the human Ndc80C construct. Regions resolved in the cryo-EM maps are shaded (NDC80: 16-24, 68-327; NUF2: 1-216). **e**, SDS-PAGE gel of the Ndc80C sample after gel filtration. Bands corresponding to the four Ndc80C subunits are labeled. **f**, Mass photometry analysis of Ndc80C shows a major peak at the expected molecular mass for the Ndc80 complex. Measured molecular masses, corresponding errors, and population percentages are indicated.

**Extended Data Fig. 2:**
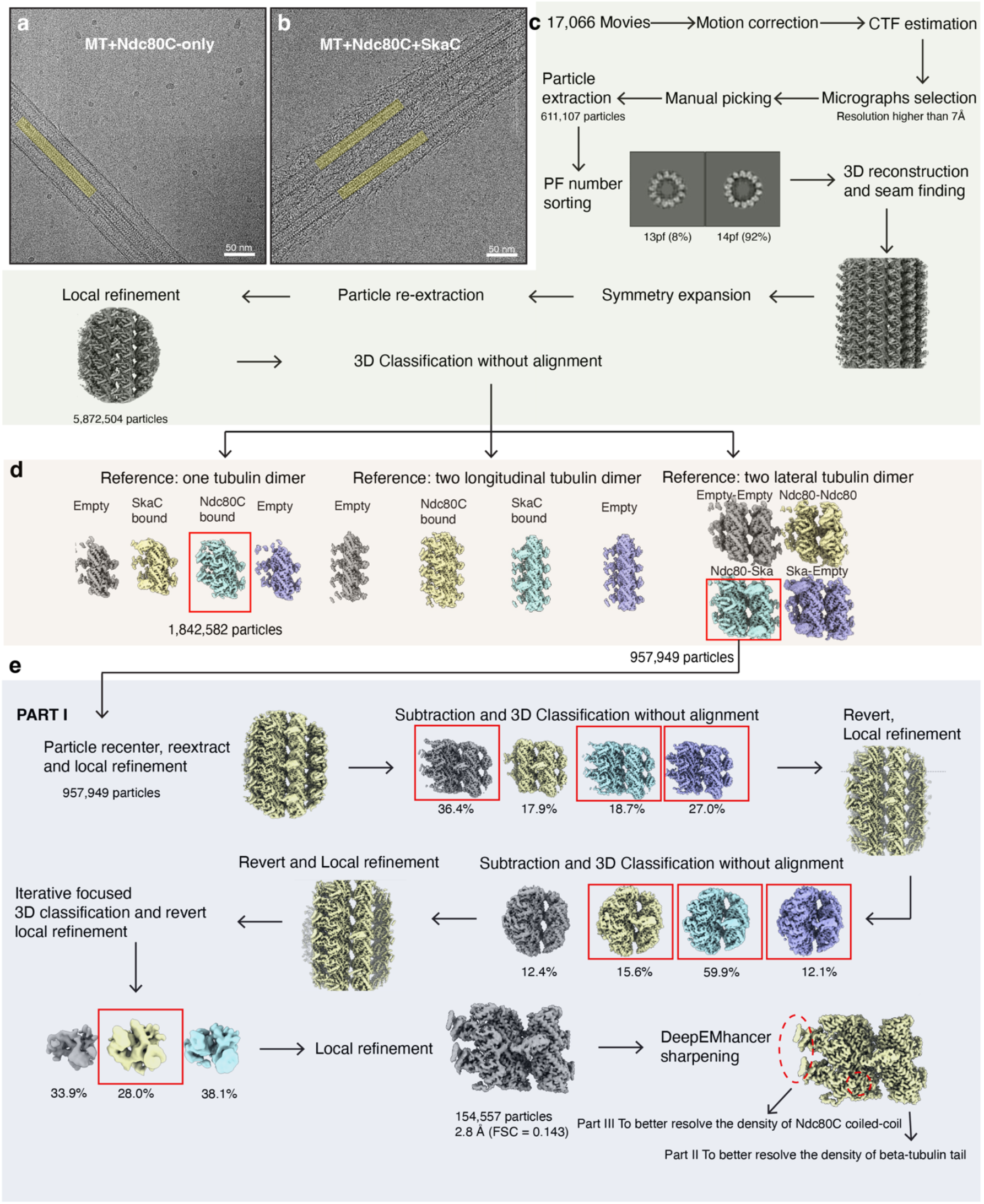
Data processing workflow for the overall MT-Ndc80C-SkaC and Tubulin-Ndc80C-SkaC subregions. a, Representative cryo-EM micrograph of the MT-Ndc80C-only sample. The MT density is highlighted in yellow (same for (**b**)). **b**, Representative cryo-EM micrograph of the MT-Ndc80C-SkaC sample. **c**, Data processing flowchart for the initial reconstruction of the MT-Ndc80C-SkaC. **d**, Focused 3D classification using masks encompassing a single tubulin dimer (left), two longitudinal tubulin dimers (middle), or two lateral tubulin dimers (right). **e**, Detailed processing flowchart for the Tubulin-Ndc80C-SkaC sub-region (Part I) analysis.

**Extended Data Fig. 3:**
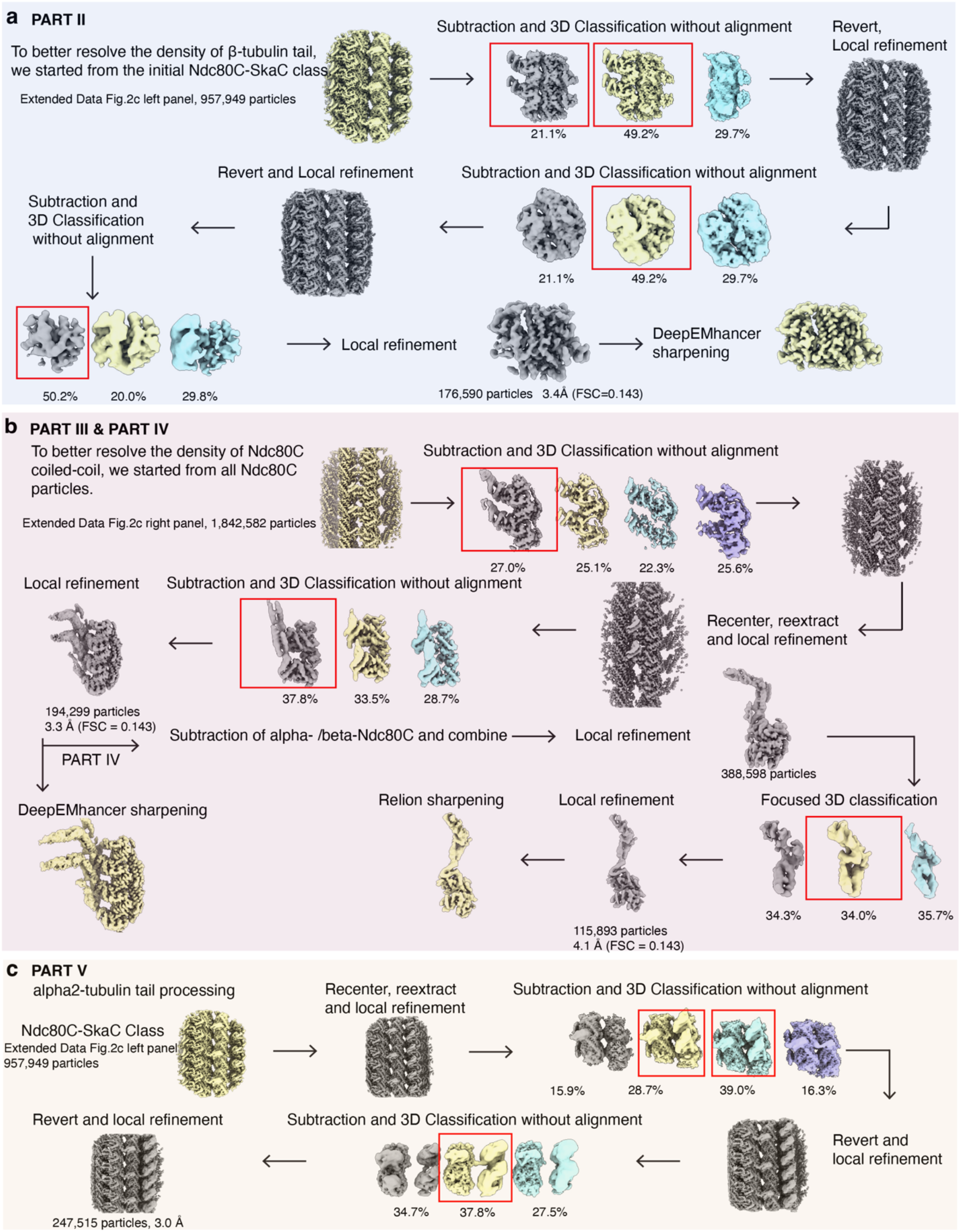
Data processing workflow of β-tubulin tail, Ndc80C coiled-coil, Ndc80C kink-SKA3 and α2-tubulin tail subregion analyses. a, Data processing flowchart for the β-tubulin tail sub-region (Part II). **b**, Data processing flowchart of two adjacent Ndc80C coiled coils (Part III) and the Ndc80C kink region with SKA3 (Part IV). **c**, Data processing flowchart for the α2-tubulin tail sub-region (Part V).

**Extended Data Fig. 4:**
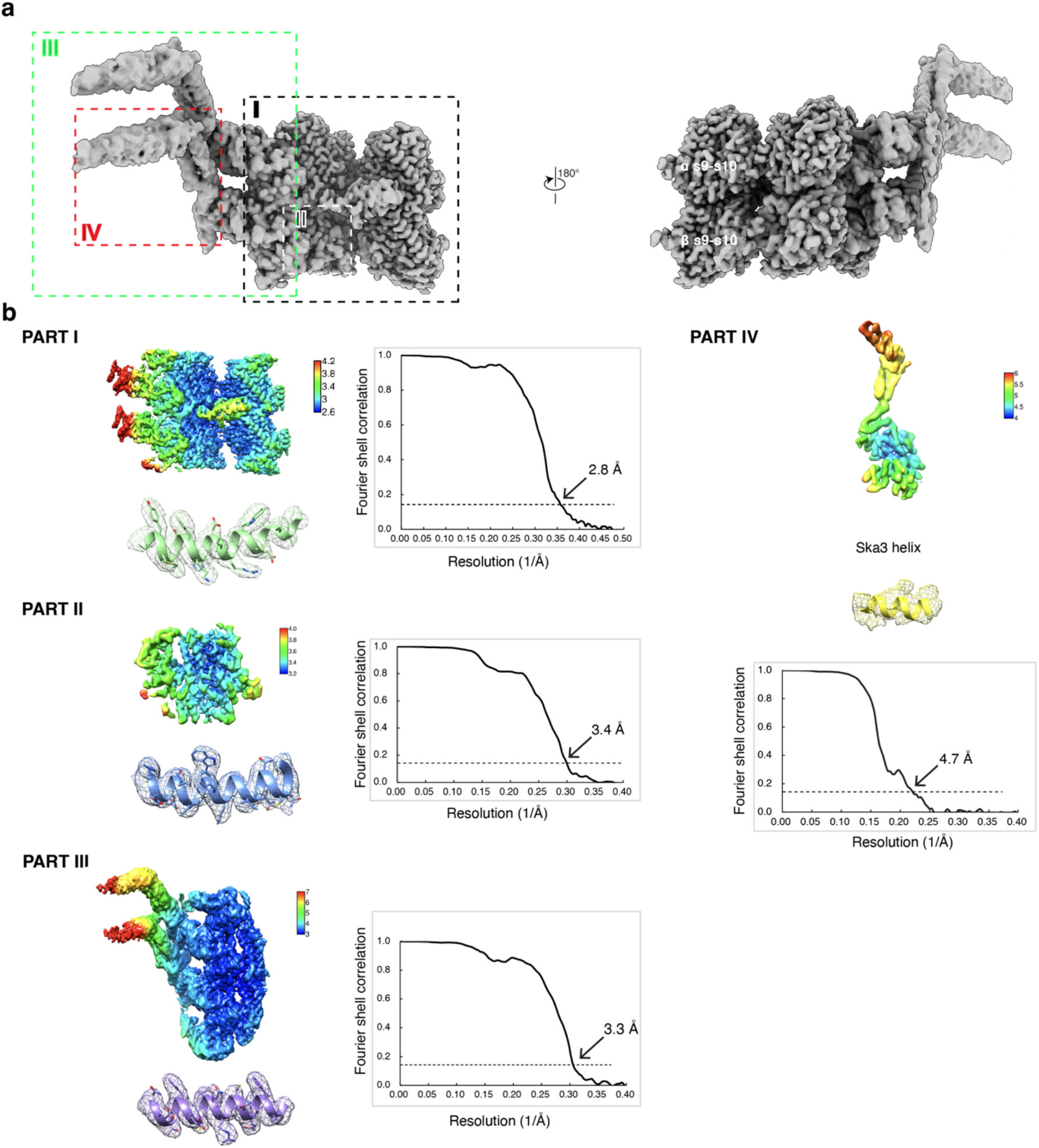
Map quality analysis for each reconstruction. a, Composite map of the Tubulin-Ndc80C-SkaC. Dashed boxes indicate regions subjected to local refinement during processing. The map clearly distinguishes the s9-s10 loop differences between α- and β-tubulin. **b**, Local-resolution maps and corresponding FSC curves for each locally refined subregion: the consensus Tubulin-Ndc80C-SkaC map (Part I) and focused maps of β-tubulin (Part II), two adjacent Ndc80Cs (Part III), and the Ndc80C kink region and SKA3 (Par IV).

**Extended Data Fig. 5:**
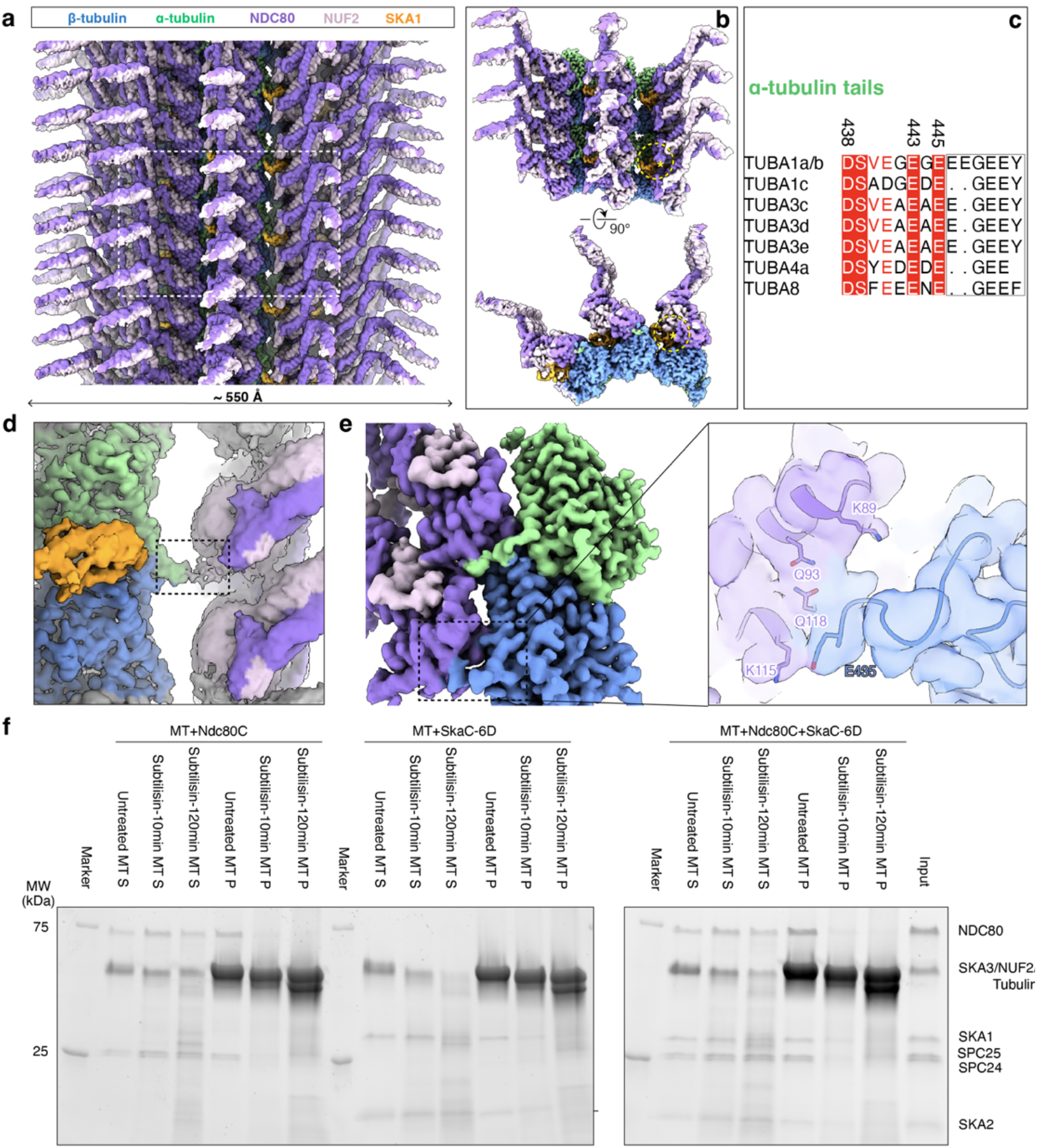
Composite, average map and tubulin-tail analyses of Ndc80C and SkaC binding on the MT lattice. a, Composite averaged cryo-EM density map of Ndc80C and SkaC bound to the MT. The dashed box marks three PFs, each containing two longitudinal tubulin dimers. **b**, Enlarged views of the boxed region in (**a**). In the averaged map, Ndc80C and SkaC densities overlap on the MT surface (asterisk, yellow dashed circle). **c**, Alignment of α-tubulin tail sequences from the indicated isoforms. **d**, Surface representation of the cryo-EM map showing the α2-tubulin tail extending toward the NDC80 N-tail on the adjacent PF. **e**, Surface representation of the cryo-EM map (left) and transparent map with the fitted atomic model (right) showing β-tubulin tail density resolved to E435 in the presence of Ndc80C. **f**, SDS-PAGE analysis of MT pelleting assays showing Ndc80C and SkaC-6D binding to untreated MTs or to MTs treated with subtilisin for 10 min or 120 min, as indicated. Pellet (P) and supernatant (S) fractions are shown. From left to right, sample groups correspond to Ndc80C alone, SkaC-6D alone, and Ndc80C plus SkaC-6D.

**Extended Data Fig. 6:**
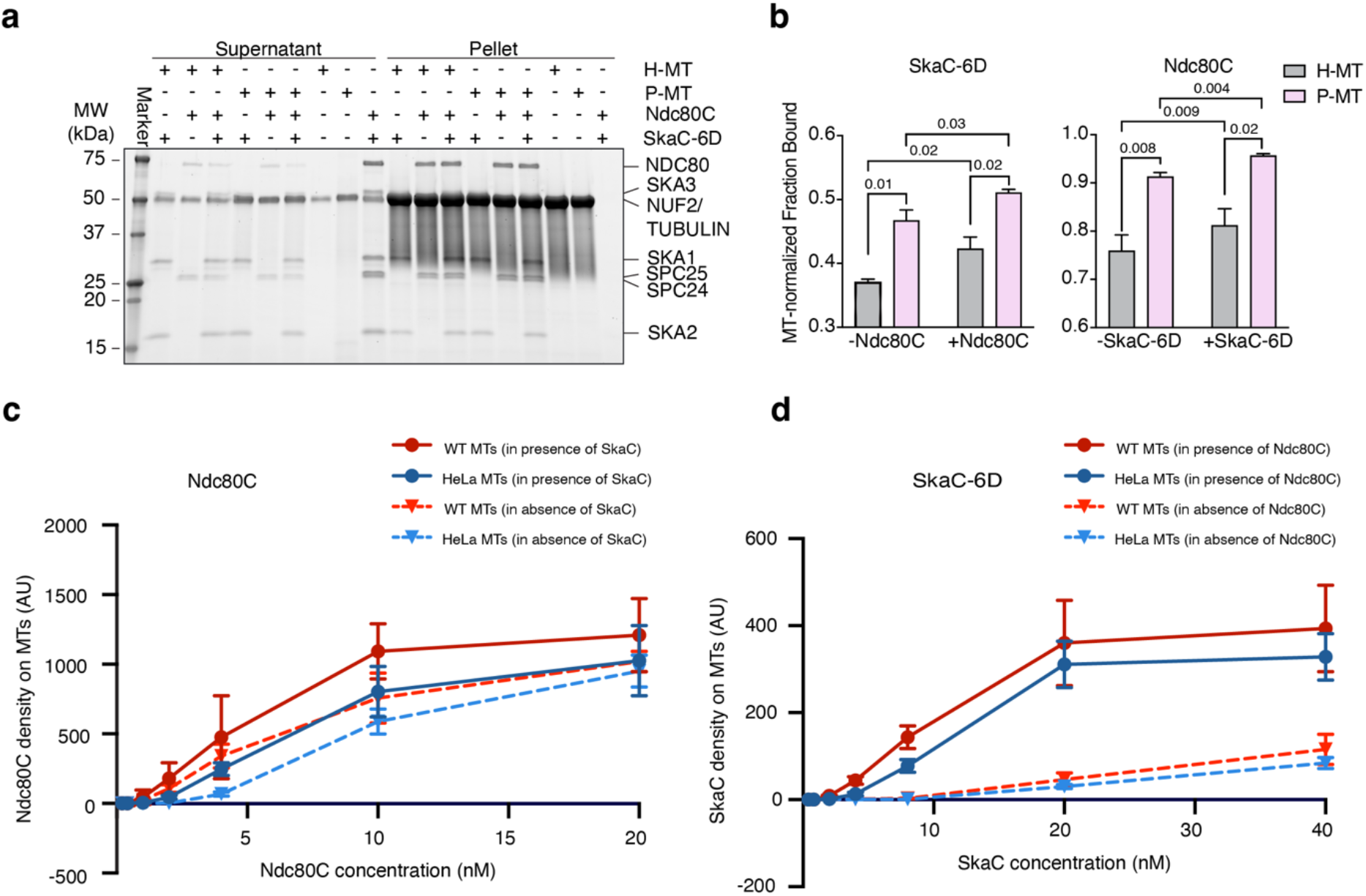
Concentration dependence of Ndc80C and SkaC binding to porcine brain and HeLa MTs. a, SDS-PAGE analysis of MT-pelleting assays shows binding of Ndc80C and SkaC to HeLa-MTs, which lack most tubulin PTMs, and to porcine brain MTs, which are enriched in PTMs. **b**, Quantification of the experiments shown in (**a**) for SkaC (left) and Ndc80C (right), with the pelleted SkaC or Ndc80C signal normalized to the corresponding pelleted MT signal (mean ± SD; three independent experiments). SkaC binding was quantified using the SKA2 band, and Ndc80C binding was quantified using the NDC80 band. P-values were calculated by a two-tailed t-test. **c**, Density of Ndc80C on GMPCPP MTs assembled from wild-type (WT) mouse brain (red) or HeLa (blue) tubulin in presence (solid lines) or absence (dotted lines) of SkaC. Data are presented as mean ± SD (n=32, 39, 43, 47 MTs, respectively in 1 independent experiment for each condition). **d**, Density of SkaC on GMPCPP MTs assembled from wild-type (WT) mouse brain (red) or HeLa (blue) tubulin in the presence (solid lines) or absence (dotted lines) of Ndc80C. Data are presented as mean ± SD (n=31, 39, 36, 36 MTs, respectively, in one independent experiment for each condition).

**Extended Data Fig. 7:**
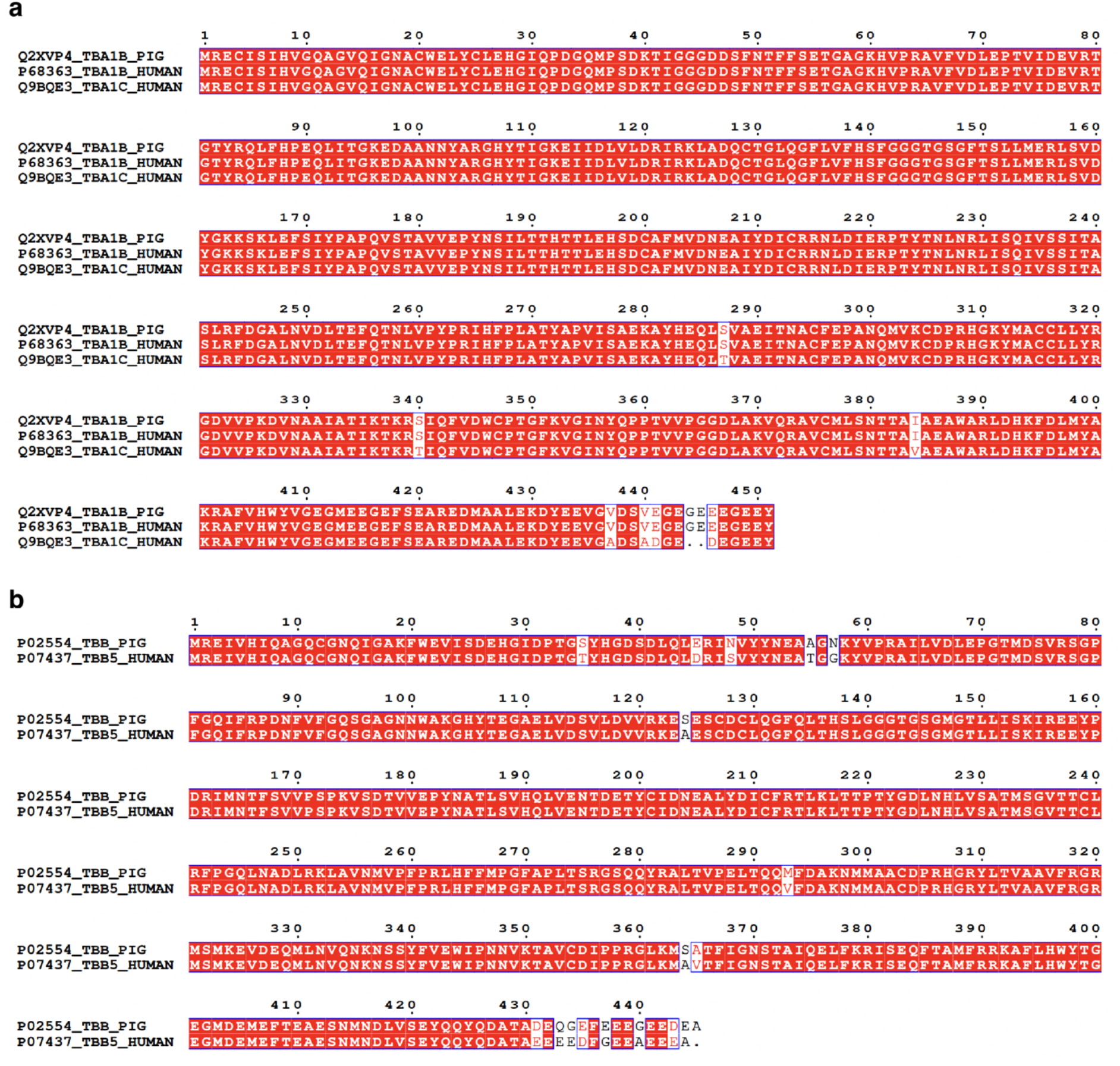
Sequence alignments between Hela and Porcine tubulin. Sequence alignments of α-(**a**) and β-tubulin (**b**) from HeLa cells and porcine brain tubulin, using the major isotype in each case (human α-tubulin: TUBA1B, UniProt P68363, and TUBA1C, UniProt Q9BQE3; human β-tubulin: TUBB/TUBB5, UniProt P07437; porcine α-tubulin: TUBA1B, UniProt Q2XVP4; porcine β-tubulin: UniProt P02554). Sequence alignments were generated with ClustalW^100^ and displayed using ESPRIPT^101^ notation, with identical residues highlighted in red and similar residues shown in red text boxed in blue. The same notation is used throughout all sequence alignments in this study.

**Extended Data Fig. 8:**
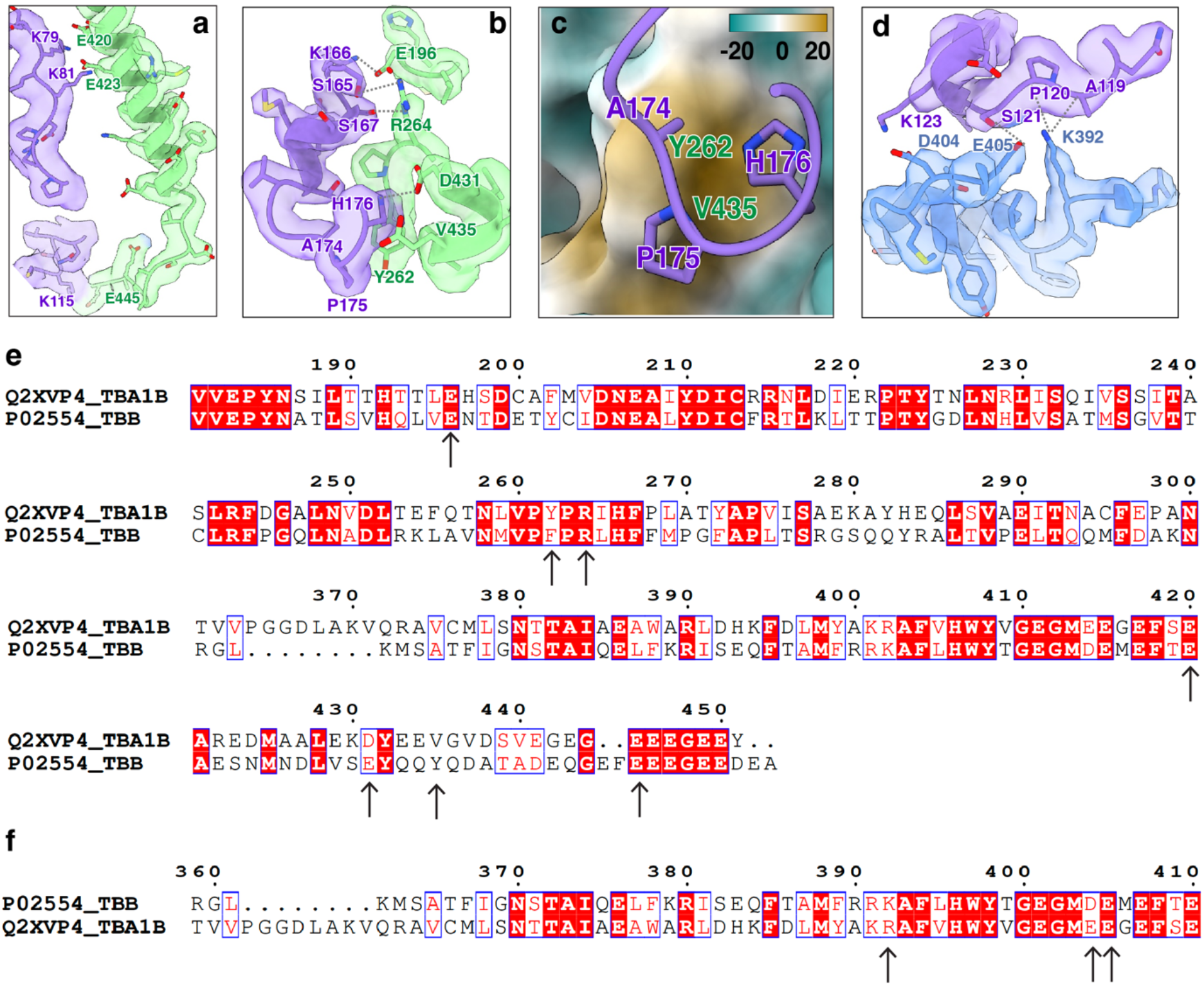
Structural and sequence conservation of the Ndc80C-MT interface. a,b, Cryo-EM density fitted with the atomic model showing the interface between α-Ndc80C and α-tubulin. Key residues are labeled, and putative salt bridges and hydrogen bonds are indicated by black dashed lines (same for (**d**)). DeepEMhancer sharpened maps are used for visualization (same for (**d**)). **c**, Hydrophobic contacts in (**b)**, with α-tubulin shown as a surface colored by the Kyte-Doolittle scale (brown, hydrophobic; green, hydrophilic; white, intermediate). α-Ndc80C shown as cartoon. The same coloring scheme is used for all hydrophobic-interaction panels. **d**, Cryo-EM map fitted with the atomic model showing the interface between α-NDC80 and β-tubulin. **e,f**, Sequence alignments of porcine α- and β-tubulin regions that contribute to the Ndc80C-binding interface. α-tubulin is used as the reference in (**e**), and β-tubulin is used as the reference in (**f**). Arrows mark residues involved in Ndc80C contacts. Several interface residues are conserved between α- and β-tubulin, while others show conservative acidic, basic, or hydrophobic/aromatic substitutions. The major porcine brain tubulin isotypes were used.

**Extended Data Fig. 9:**
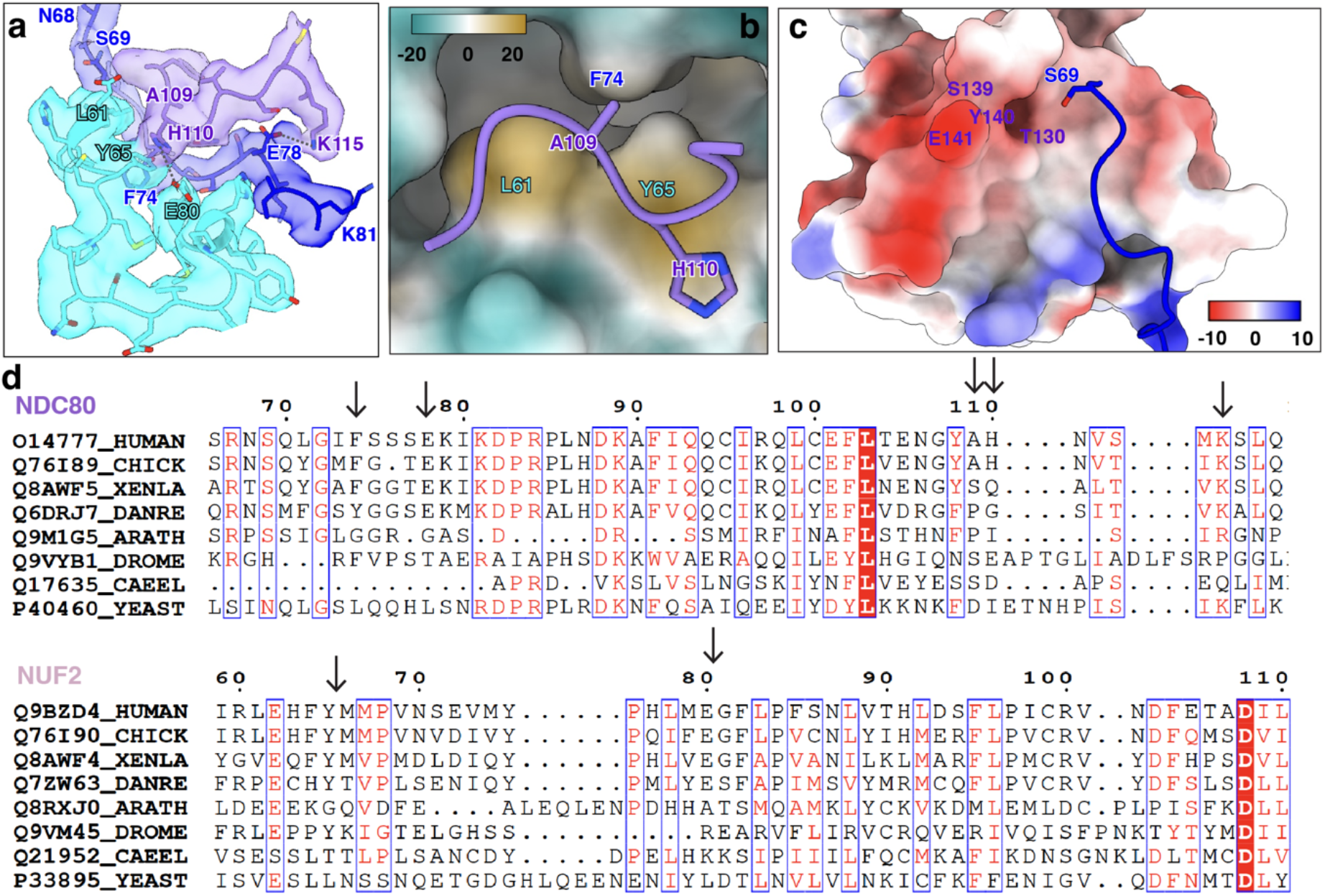
Structural and evolutionary features of a putative cooperative interface between neighboring Ndc80 complexes. a, Cryo-EM map fitted with model showing the interface between α-NDC80 and β-NDC80. **b**, Hydrophobic contacts in (**a**). **c**, Electrostatic surface of the α-Ndc80C and cartoon of β-Ndc80C showing NDC80 S69 facing an acidic surface in the former. **d**, Sequence alignment of NDC80 residues 66-118 and NUF2 residues 59-110. Residues contributing to cooperative binding are indicated by black arrows. Residue K115 in NDC80 is largely conserved across species, except in Drosophila and C. elegans. In contrast, NDC80 residues F74 and E78, as well as NUF2 Y65 and E80, are highly conserved but only among vertebrates. By comparison, NDC80 A109/H110 and NUF2 L61 show limited conservation within vertebrates. These conservation patterns indicate that the cooperative binding interface is selectively reinforced in vertebrates. Sequence labels denote UniProt accession and species: HUMAN (*Homo sapiens*); CHICK (*Gallus gallus*); XENLA (*Xenopus laevis*); DANRE (*Danio rerio*); ARATH (*Arabidopsis thaliana*); DROME (*Drosophila melanogaster*); CAEEL (*Caenorhabditis elegans*); YEAST (*Saccharomyces cerevisiae*).

**Extended Data Fig. 10:**
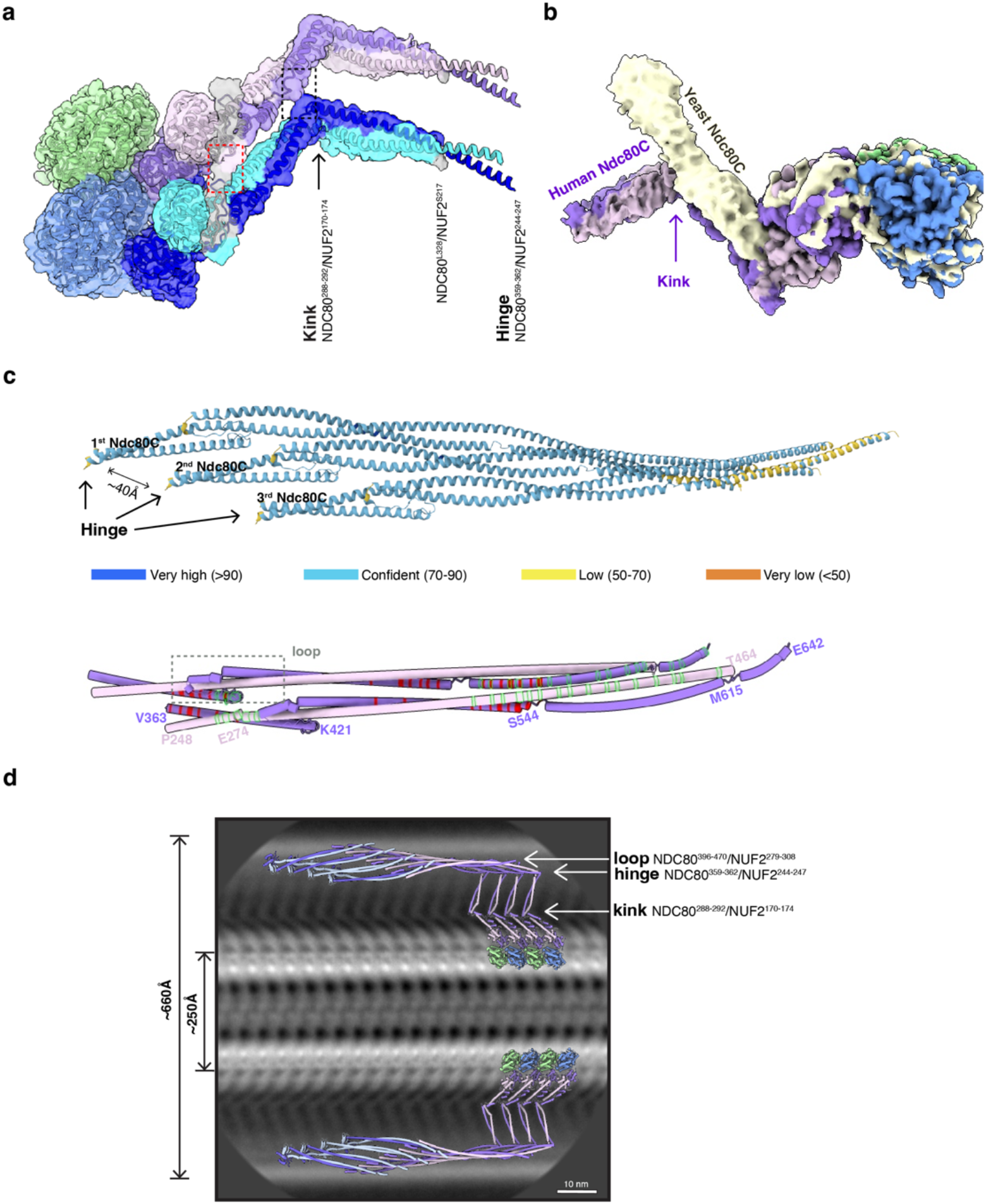
Additional regions of the Ndc80C coiled-coil that may contribute to cooperativity, and the composite full-length model of human Ndc80C. a, Cryo-EM map showing two Ndc80C bound to a single tubulin dimer. The black dashed box highlights the region of possible contact between adjacent Ndc80C at the kink; the red dashed box indicates an additional region between adjacent tails. The kink, canonical hinge, and terminal residues of our model are listed and marked by black arrows. **b**, Superposition of human (this study) and yeast NDC80 and NUF2 (EMDB: 18304) cryo-EM maps highlights a kink in the human Ndc80C coiled-coil region that is absent in yeast (beige). **c**, AF3-predicted model of three Ndc80C coiled-coil regions (NDC80^367-642^/NUF2^250-464^) arranged as a bundle with ∼40 Å spacing between adjacent coiled-coils (upper panel), colored based on confidence of the prediction. For clarity, the lower panel shows only two AF3 models, with residues contributing to intermolecular contacts highlighted in red and green. The dashed box marks the loop region. **d**, A representative 2D class average from the MT-Ndc80C-only sample. The composite full-length of human Ndc80C model using experimental section and AF3 for the rest of the coiled-coil is overall consistent with the observed density in 2D average. Kink, canonical hinge and loop regions are indicated by white arrows for the first complex on the top right.

**Extended Data Fig. 11:**
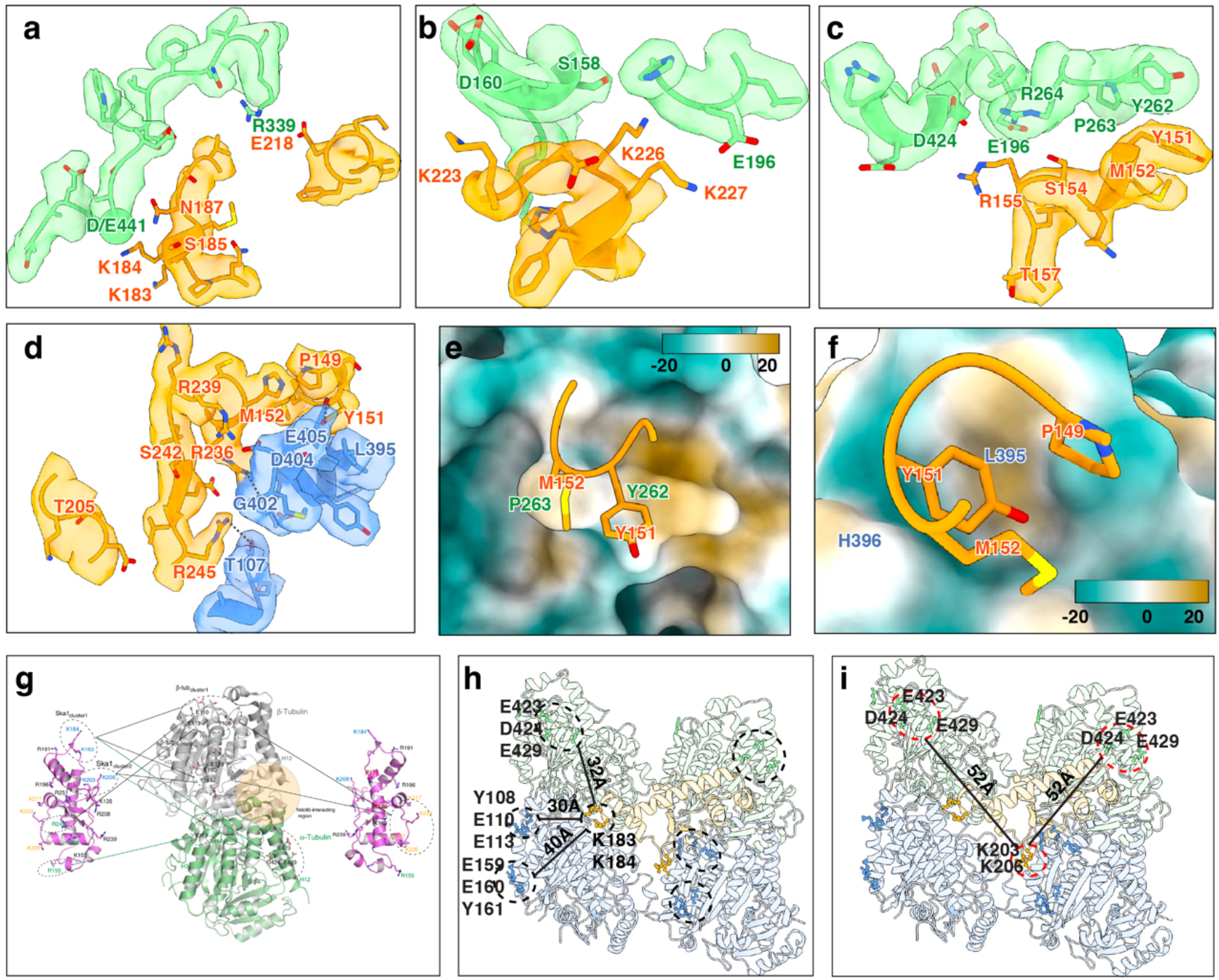
Structural basis of SKA1-MTBD binding to the MT lattice and comparison with XL-MS interaction sites. **a**-**d**, Cryo-EM density fitted with atomic models showing the interface between SKA1-MTBD and α1-tubulin (**a**), α2-tubulin (**b**-**c**), and β2-tubulin (**d**). Key residues are labeled and putative salt bridges and hydrogen bonds are indicated by black dashed lines. DeepEMhancer sharpened maps are used for visualization (same for (**a**-**d**)). **e**-**f**, Hydrophobic contacts between SKA1 and α2-tubulin (**e**) or β2-tubulin (**f**), with SKA1 shown as a cartoon and tubulin rendered as hydrophobic/hydrophilic surfaces using the Kyte-Doolittle coloring. **g**, Previously published XL-MS data^33^ identifying interaction patches between SKA1-MTBD and tubulin, reproduced from Abad et al. (2014) under a CC BY 3.0 license. **h,i**, For XL-MS patches involving SKA1 K183/K184 (**h**) or K203/K206 (**i**), the corresponding positions in our structure are separated by more than 30 Å, suggesting that these contacts are not captured by the straight-lattice interface resolved here and may instead arise from interactions with curved tubulin conformations.

**Extended Data Fig. 12:**
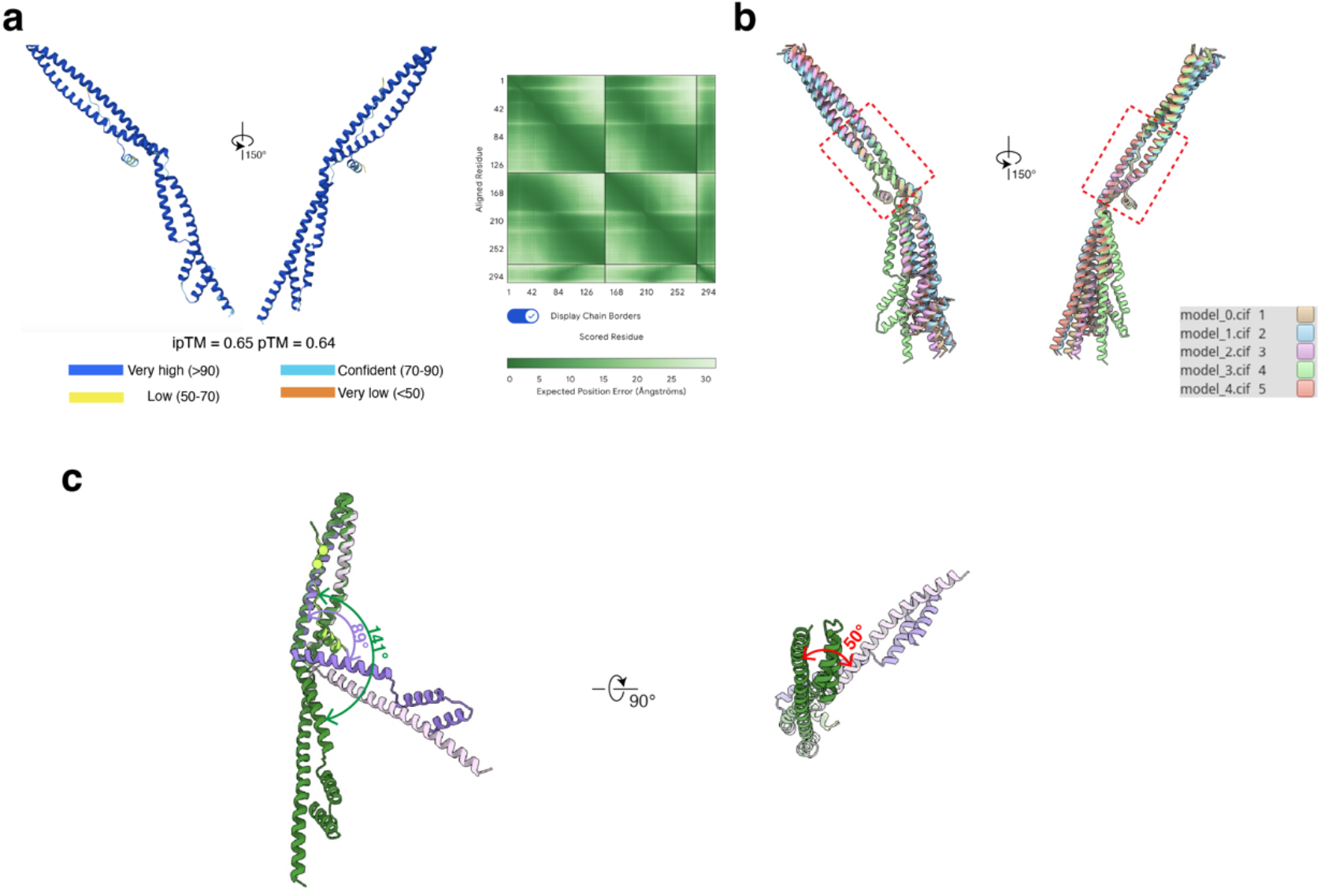
Comparison of the AF3-predicted structure with our cryo-EM based model for the Ndc80C coiled-coil kink region interacting with SKA3. a, AF3 predicted structure for NDC80^223-360^/NUF2^116-245^/SKA3-6D^355-380^ colored based on model confidence. The corresponding predicted aligned error (PAE) plot is shown on the right. **b**, All five AF3 models from panel (**a**) are superimposed on the SKA3 tether helix and show differences in the angle at the kink. The red dashed box highlights the conserved interaction interface common to all models. **c**, Superposition of our experimentally based model (colored as in the main figures) with the highest-confidence AF3 model (shown in green) shows a similar interaction interface but a distinct Ndc80C kink angle.

**Supplemental Data Table 1.**
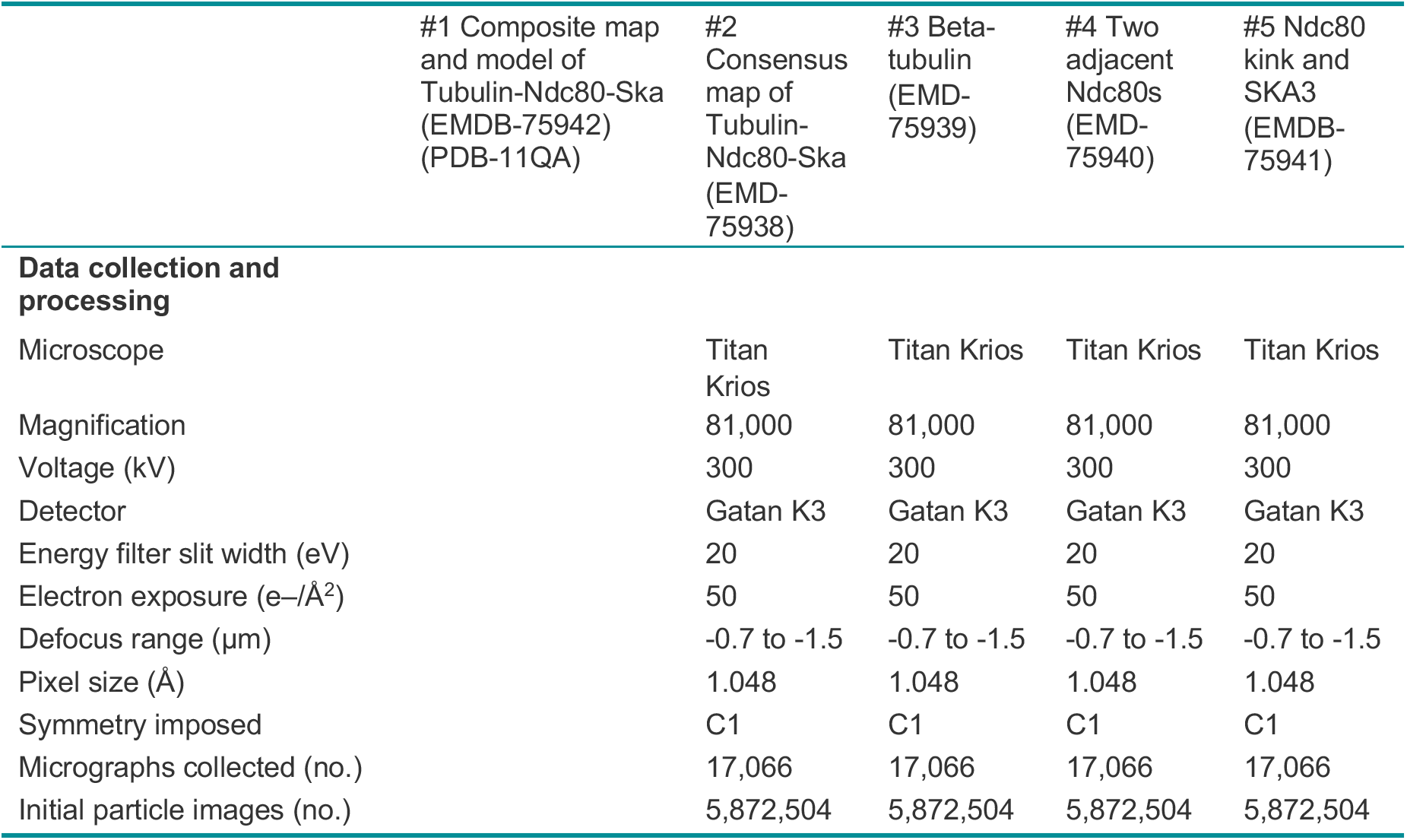
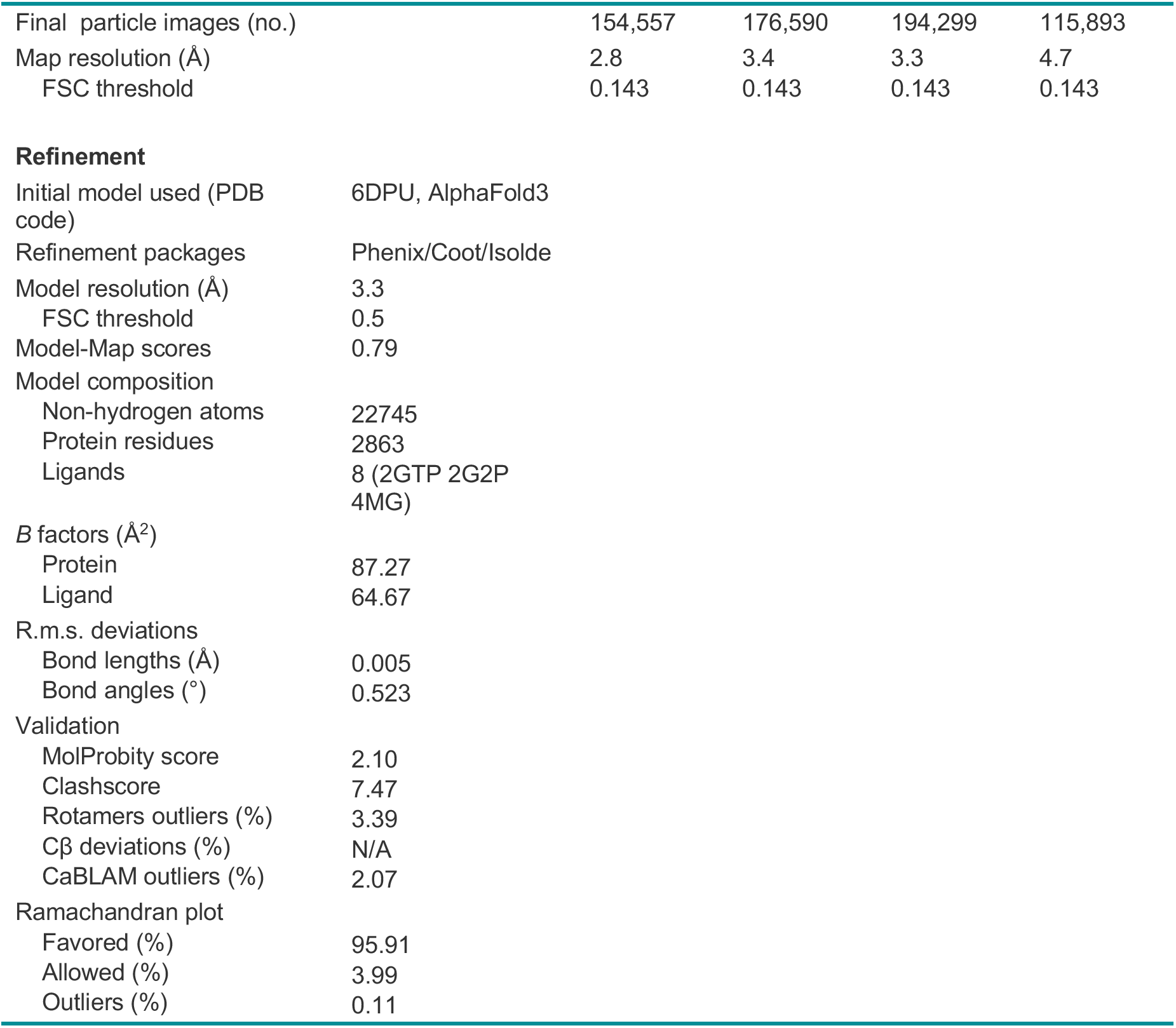
Cryo-EM data collection, refinement and validation statistics.

